# Targeted single-nucleus sequencing of 39,800 neurons reveals extensive low-frequency somatic variants

**DOI:** 10.64898/2026.09.21.753260

**Authors:** Vanshika Bidhan, Sarah Wynants, Toon Swings, Cristina T. Vicente, Fahri Küçükali, Marleen Van den Broeck, Jeroen GJ. van Rooij, Merel O. Mol, Laura L. Donker Kaat, Safa Al-Sarraj, Istvan Bodi, Andrew King, Claire Troakes, Christos Proukakis, Jolien Schaeverbeke, Dietmar R. Thal, Rik Vandenberghe, Mathieu Vandenbulcke, Aivi T. Nguyen, R. Ross Reichard, Julia Kofler, Oscar L. Lopez, Charles L. White, Bradley F. Boeve, Neill R. Graff-Radford, Keith A. Josephs, Ronald C. Petersen, Nikhil B. Ghayal, Melissa E. Murray, Dennis W. Dickson, John C. van Swieten, Kristel Sleegers, Harro Seelaar, Wouter De Coster, Rosa Rademakers

**Affiliations:** Department of Biomedical Sciences, University of Antwerp, Antwerp 2610, Belgium; VIB Center for Molecular Neurology, VIB, Antwerp 2610, Belgium; VIB Technology Watch, Technology Innovation Lab, VIB, Leuven 3000, Belgium; Department of Neurology, Erasmus Medical Center, Rotterdam 3015 CN, The Netherlands; Department of Clinical Genetics, Erasmus Medical Center, Rotterdam 3015 CN, The Netherlands; London Neurodegenerative Diseases Brain Bank, Department of Basic and Clinical Neuroscience, Institute of Psychiatry, Psychology & Neuroscience, King’s College London, London SE5 8AF, UK; Department of Clinical Neuropathology, King’s College Hospital NHS Foundation Trust, London SE5 9RS, UK; Department of Clinical and Movement Neurosciences, UCL Queen Square Institute of Neurology, London NW3 2PF, UK; Laboratory for Cognitive Neurology, Department of Neurosciences, Leuven Brain Institute, KU Leuven, Leuven 3000, Belgium; Laboratory of Neuropathology, Department of Imaging and Pathology, and Leuven Brain Institute, KU Leuven, Leuven 3000, Belgium; Department of Pathology, University Hospital Leuven (UZ Leuven), Leuven 3000, Belgium; Department of Neurology, University Hospitals Leuven (UZ Leuven), Leuven 3000, Belgium; Neuropsychiatry, Department of Neurosciences, Leuven Brain Institute, KU Leuven, Leuven 3000, Belgium; Department of Geriatric Psychiatry, University Hospitals Leuven (UZ Leuven), Leuven 3000, Belgium; Department of Laboratory Medicine and Pathology, Mayo Clinic, Rochester MN, 55905, USA; Department of Pathology, University of Pittsburgh, Pittsburgh, PA 15213, USA; Department of Neurology, University of Pittsburgh, Pittsburgh, PA 15213, USA; Division of Neuropathology, University of Texas Southwestern Medical Center, Dallas, TX 75390, USA; Department of Neurology, Mayo Clinic, Rochester, MN 55905, USA; Department of Neurology, Mayo Clinic, Jacksonville, FL 32224, USA; Department of Neuroscience, Mayo Clinic, Jacksonville, FL 32224, USA; Department of Laboratory Medicine and Pathology, Mayo Clinic, Jacksonville, FL 32224, USA

**Keywords:** Somatic mutations, Neuronal somatic mosaicism, Frontotemporal dementia, *TARDBP*

## Abstract

Recent evidence implicates somatic mutations in disease-associated genes across multiple neurodegenerative disorders, including the identification of two somatic *TARDBP* variants as a putative cause of frontotemporal lobar degeneration with TDP-43 pathology type C (FTLD-TDP type C). However, these findings remain anecdotal.

We performed single-nucleus amplicon sequencing of 39,800 neurons from the superior temporal gyrus of sporadic FTLD-TDP type C patients (34,738 neurons, 52 individuals) and non-demented controls (5,062 neurons, 26 individuals) using the Mission Bio Tapestri platform and investigated somatic variants in *TARDBP* and other FTLD-TDP-associated genes in a case-control setting.

We uncovered an extensive landscape of ultra-low-frequency (<1%) somatic mutations across all analysed genes, including *TARDBP*. Rare somatic occurrences of known ALS/FTD-associated germline *TARDBP* variants, including p.N267S and p.M337V, were identified exclusively in patients. Among all targeted genes, *TARDBP* had the highest proportion of neurons harbouring a somatic variant, and this mutational burden was lower in individuals who died at an older age. Strikingly, the C-terminal domain of TDP-43, where most ALS/FTD germline pathogenic variants reside, exhibited a lower neuronal mutational burden than its other domains. This suggests that neurons harbouring damaging mutations may undergo progressive loss prior to autopsy, leaving only rare survivors at frequencies too low to detect by bulk or small-scale single-cell approaches. Together, our findings provide an unprecedented resolution into the somatic mutational landscape of post-mitotic neurons in FTLD-TDP type C and reveal a gene-specific layer of neuronal mosaicism that current single-cell WGS studies fail to detect.

## Introduction

Frontotemporal dementia (FTD) is a common form of early-onset dementia, marked by changes in behaviour, language or motor function. FTD is most often caused by an underlying frontotemporal lobar degeneration (FTLD), with subtypes defined based on the aggregating proteins. FTLD with neuronal and glial cytoplasmic aggregates of TAR DNA-binding protein 43 (FTLD-TDP) is the largest neuropathological subgroup,^1–5^ and is classified into neuropathological subtypes (A–E) based on the morphology and distribution of TDP-43 inclusions. FTLD-TDP type C is characterized by long dystrophic neurites and cytoplasmic inclusions that predominantly affect the (sub)cortical regions of the anterior temporal lobe.^6–9^ These inclusions were recently resolved as heteromeric amyloid fibrils in which TDP-43 co-assembles with annexin A11.^10,11^ Clinically, patients typically present with semantic variant primary progressive aphasia, although behavioural-predominant presentations associated with right-sided or more distributed anterior temporal involvement also occur.^8,9,12–14^ FTLD-TDP type C patients are largely without a family history of FTLD, with no germline monogenic cause established to date, though rare variants have been associated with this subtype.^15,16^

Somatic mutations in the brain have long been hypothesized as a contributor to sporadic neurodegenerative diseases, and emerging evidence now implicates somatic variants in genes otherwise known to cause familial disease. Deep targeted sequencing of ALS (Amyotrophic Lateral Sclerosis) and FTD-associated genes in two independent studies has revealed an enrichment of low-level somatic variants in sporadic ALS and FTD patients relative to controls,^17,18^ with deleterious somatic mutations identified in ∼2% of sporadic ALS and FTD patients.^17^ In a whole-exome deep sequencing study investigating somatic variation in a cohort of 16 FTLD-TDP type C patients, a rare somatic *TARDBP* p.R42H variant was identified in the middle temporal gyrus (MTG) of one patient (1.4%) and was additionally observed in the parietal (1.2%) and frontal lobes (0.5%) of the same carrier. The authors also identified a somatic *TARDBP* p.L41F variant in a second patient (2%).^19^ Unlike most pathogenic germline *TARDBP* mutations in ALS and FTD that affect the low-complexity C-terminal domain (CTD), these two somatic variants affect the N-terminal domain (NTD) and may promote TDP-43 aggregation through N-terminal misfolding.^19^ These findings implicated somatic *TARDBP* mutations as a potential contributing factor to FTLD-TDP type C. The absence of relevant mutations in the remaining 14 cases was attributed to limited sensitivity, stringent filtering, neuronal loss, sampling bias, possible somatic variants in other TDP-43-related genes or other non-genetic causes. Moreover, rare observations of somatic variants in non-demented individuals were attributed to potential sequencing errors.^19^

Building on these findings and limitations, we performed single-nucleus amplicon sequencing of neurons to investigate whether somatic *TARDBP* mutations could partially explain the sporadic FTLD-TDP type C pathology in a large cohort of affected individuals. To our knowledge, this represents the first targeted single-nucleus analysis of somatic mutations in FTLD-TDP and the first application of single-nucleus amplicon sequencing using the Tapestri platform in a neurodegenerative disorder. To reduce bias introduced by neuronal loss, we analysed superior temporal gyrus (STG) samples, a region characterized by moderate-to-severe pathology but with mild-to-moderate neuronal loss in FTLD-TDP type C patients.^20^ Beyond *TARDBP*, we also analysed somatic mutations in six other FTLD-TDP-associated genes.

Previous single-cell studies investigating brain somatic mutations in neurodegenerative diseases have used single-cell whole-genome sequencing (scWGS), which has been limited to fewer than 500 neurons in total.^21–25^ In our study, we assessed somatic mutations in 39,800 STG neurons from 52 FTLD-TDP type C patients and 26 neuropathologically normal controls and confirmed the previously reported *TARDBP* p.R42H somatic variant in the known carrier. Surprisingly, we identified 19,102 coding somatic single-nucleotide variants (sSNVs) and small indels across the targeted regions, with >90% of variants detected in only a single neuron within a sample. While potentially pathogenic somatic *TARDBP* variants affecting the NTD were detected in multiple patients, variants in its CTD were significantly underrepresented, possibly due to survivor bias resulting from selective loss of neurons harbouring relevant variants. However, we still identified patient-exclusive *TARDBP* sSNVs that correspond to known pathogenic germline variants implicated in ALS and FTD, including p.N267S and p.M337V. Our findings underscore the need for large-scale approaches at single-cell resolution to detect potentially pathogenic brain somatic mutations, as their low allele frequencies make them largely undetectable by other methods.

## Materials and methods

### Cohort selection

This study included 96 frozen brain tissue samples from 95 different individuals. We included 93 samples from the STG of 57 FTLD-TDP type C patients and 36 neuropathologically normal controls from seven collaborating sites **(Supplementary Table 1** and **Supplementary material, ‘Materials and methods’ section**). These samples were selected based on the neuropathological diagnoses made at each site. From the patient carrying the *TARDBP* p.R42H somatic variant, we further included an MTG sample (N1_P, positive control) in addition to the STG sample (T1_P) to confirm the originally reported frequency.^19^ The two remaining samples were frontal cortex-derived (one patient and one control, unrelated to the STG cohort) and were included for a pilot comparison of somatic mutation frequencies between unsorted and NeuN+-sorted nuclei (**Supplementary material, ‘Pilot experiment’ section**). Somatic variants from these two samples were excluded from downstream analyses. All 95 individuals provided informed consent for their samples to be used in research.

### Amplicon panel design

We designed a custom amplicon panel targeting exons of known FTLD-TDP-associated genes, common germline SNPs for sample demultiplexing and amplicons spanning chromosome Y (**Supplementary Tables 2and 3** and **Supplementary material, ‘Materials and methods’ section**). In addition to *TARDBP*, we included *GRN*, *OPTN*, *TBK1*, *TMEM106B* and *UNC13A* to investigate if somatic variation in these genes could contribute to FTLD-TDP type C and to compare their somatic mutational burden with that of *TARDBP*. We also included *TET2*, as an enrichment of somatic *TET2* variants has been reported in blood from FTD patients,^26,27^ possibly confounded by clonal hematopoiesis.^28,29^

### Targeted single-nucleus amplicon sequencing

We adopted a pooled-sample design, combining five tissue samples, weighing 30 mg each, per pool (with the exception of sample O1_P). Nuclei were isolated and enriched using the MARS system (Applied Cells), and NeuN+ nuclei were subsequently isolated using fluorescence-activated nuclei sorting (FANS). DNA libraries were prepared on the Mission Bio Tapestri platform (v2 chemistry) and sequenced on an Element Biosciences AVITI sequencer (**Supplementary material, ‘Materials and methods’ section**). Across all pools, we recovered 47,200 nuclei at a mean depth of 463 reads per nucleus per amplicon (range: 64–1,798) prior to variant filtering (**Supplementary Table 4**). Samples were deconvoluted using a germline SNP-based approach, and doublets were identified with an artificial doublet strategy (**Supplementary material, ‘Materials and methods’ section**). The full workflow is summarized in **Fig. 1A** and **B**. Raw sequencing reads were processed using Mission Bio’s Tapestri analysis pipeline (v2.0.2), and the resulting output was analysed using Mission Bio’s mosaic pipeline (v3.12, https://missionbio.github.io/mosaic/).

**Figure 1.**
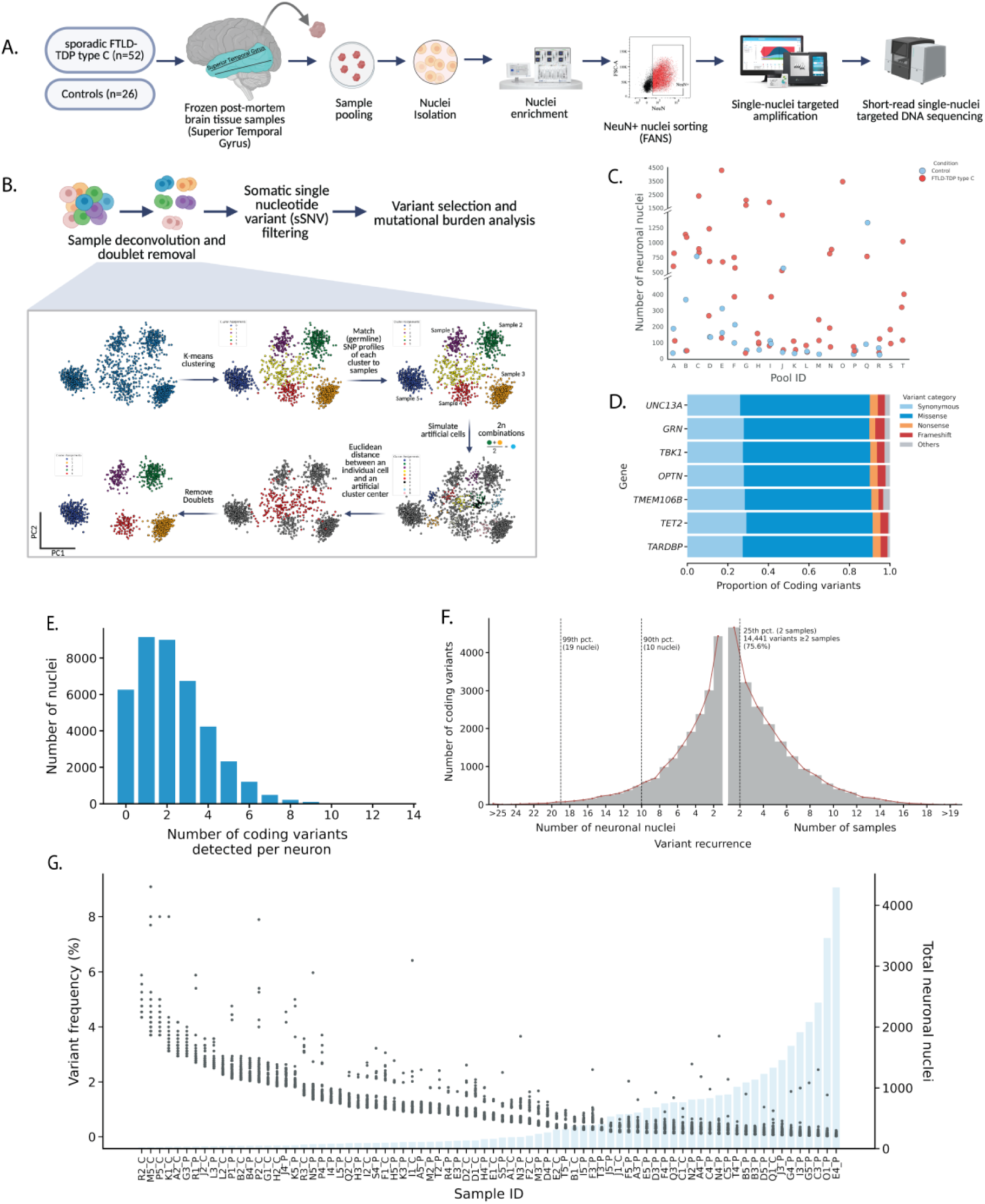
Study design, data processing and overview of somatic variants detected across FTLD-TDP-associated genes in single neurons from FTLD-TDP type C patients and neurologically healthy controls. **(A)** Superior temporal gyrus (STG) tissue samples from 52 FTLD-TDP type C patients and 26 controls were pooled in batches of five, followed by nuclei isolation and enrichment, and fluorescence-activated nuclei sorting (FANS) for NeuN+ neuronal nuclei. Sorted nuclei were subjected to target amplification on the Mission Bio Tapestri v2 platform, with short-read sequencing performed on the AVITI sequencer. **(B)** Raw sequencing data were processed, followed by sample deconvolution and doublet removal. Stringent quality filtering was applied to identify high-confidence somatic variants. **(C)** Number of NeuN+ neuronal nuclei profiled per sample following deconvolution and quality control (y-axis), grouped by pool identifier (x-axis). Each dot represents one sample, coloured by disease status. **(D)** Proportion of coding somatic variants (x-axis) per variant class (synonymous, missense, nonsense, frameshift and others) detected per targeted gene (y-axis) across the cohort. **(E)** Distribution of the number of coding somatic variants detected per neuron (x-axis). **(F)** Variant recurrence across nuclei and samples, depicted as the total number of coding somatic variants (y-axis) against the number of nuclei (left x-axis), and the number of samples with each variant (right x-axis). Over 75% of variants are shared across individuals, and 90% of variants are detected in ≤10 nuclei each, with fewer than 1% detected in more than 20 nuclei. **(G)** Variant allele frequency (VAF, left y-axis) and the total number of neuronal nuclei profiled (grey bars, right y-axis) are depicted per sample (x-axis). Samples with fewer than 500 nuclei show inflated VAFs compared with samples with more neuronal nuclei profiled. Schematics (A) and (B) were created in BioRender.

### Variant filtering

We applied a stringent, multi-tiered filtering strategy to identify high-confidence somatic variants (**Supplementary Figs. 1 and 2** and **Supplementary material, ‘Materials and methods’ section**). Variants were first filtered at the amplicon level based on amplicon performance, genomic context and population frequency. Sample-level filtering was then applied, excluding variants with poor callability (defined as the number of nuclei genotyped at a given position). Germline variants were subsequently identified and removed from each sample (**Supplementary Table 5**). Finally, nucleus-level filtering was performed, comprising binomial filtering to exclude false-positive heterozygous calls (**Supplementary Fig. 3**) and removal of variants located in close proximity within the same nucleus.

### Statistical analyses

We defined the mutational burden across a gene in a sample as the proportion of nuclei harbouring at least one somatic coding variant. Because the panel included only seven genes, the metric reflects the net cellular burden of somatic mutation rather than the per-cell mutational load. Covariate-adjusted inference on mutational burden was performed using a negative binomial generalized linear mixed-effects model (nbinom2) fit with the glmmTMB package in R,^30^ motivated by significant overdispersion under a Poisson assumption. The number of affected nuclei per sample was modelled as the response variable, with a log-transformed offset for the total number of nuclei per sample to account for differences in neuronal sampling across samples; disease status, age at death, sex, brain hemisphere and sequencing depth were included as covariates. For comparisons across TDP-43 domains and genes, an additional offset for domain/gene length was incorporated (see **Supplementary material, ‘Materials and methods’ section** for detailed model parametrization). Likelihood ratio tests (LRT) were used to compare nested models and assess interaction terms.

## Results

### Single-nucleus amplicon sequencing reveals a strikingly high number of low-frequency somatic variants

We sequenced 39,800 neuronal nuclei from 78 STG tissue samples, comprising 34,738 nuclei from 52 FTLD-TDP type C patients and 5,062 nuclei from 26 neuropathologically normal controls (**Fig. 1A** and **Supplementary Table 1**). We further analysed: (i) 587 nuclei from an MTG tissue sample from the carrier of a known somatic *TARDBP* p.R42H mutation^19^; and (ii) 741 nuclei from two frontal cortex tissue samples (**Supplementary material, ‘Pilot experiment’ section**). Nuclei recovery varied substantially across samples (range: 23–4,296) and was consistently lower in control individuals, with on average seven-fold fewer nuclei compared to patients within pooled runs (**Fig. 1C**). We also profiled mosaic loss of chromosome Y (LOY) in males, which showed LOY in 21 of 33 male patients and six of 12 male controls, with comparable frequencies between groups (patients: 0.3%, 71/23,464; controls: 0.31%, 11/3,570), consistent with the previously reported frequency of 0.31% in human neurons.^31^

Variants were called in intervals totalling 22,585 bp, after excluding low-quality amplicons, repetitive regions and regions with insufficient callability. We identified 28,345 somatic variants at 20,450 unique positions, collectively impacting 90.5% of the genotyped positions (**Supplementary Fig. 1**). Somatic SNVs accounted for the vast majority (*n* = 27,109; 95.6%), with fewer somatic insertions (*n* = 67; 0.2%) and deletions (*n* = 1,169; 4.2%). Somatic indels were predominantly single-base events (insertions = 59, deletions = 1,034), with very few multi-base events (*n* = 143) and only two deletions exceeding 10 bp. We observed a total of 19,102 somatic coding variants spanning 13,957 bp (**Table 1**), with median per-nucleus coverage of 118x in patients and 135x in controls (**Supplementary Fig. 4**). The proportions of synonymous, missense, nonsense and frameshift variants were consistent across genes (**Fig. 1D**).

**Table 1.** Somatic coding variants detected per targeted gene. The targeted coding interval size (bp), the number of amplicons, the median coverage per neuron and the number of unique base positions affected by a somatic variant are listed for each targeted gene. Somatic variant counts are reported per variant class: somatic single-nucleotide variants (sSNVs) and somatic insertions and deletions (sIndels). Non-synonymous variants are further classified as missense, nonsense, frameshift, start loss, stop loss and in-frame deletion. All variant counts reflect somatic variants that pass quality-control filters across all superior temporal gyrus samples in the cohort.

|  | *UNC13A* | *TET2* | *OPTN* | *TBK1* | *GRN* | *TARDBP* | *TMEM106B* |
| --- | --- | --- | --- | --- | --- | --- | --- |
| Targeted coding interval size | 4426 | 4291 | 1733 | 1729 | 1532 | 1099 | 806 |
| Number of amplicons | 40 | 26 | 13 | 17 | 11 | 7 | 7 |
| Median coverage per neuron | 65 | 125 | 97 | 107 | 31 | 92 | 62 |
| Unique base positions affected | 4029 | 3730 | 1491 | 1787 | 1133 | 1040 | 747 |
| **Number of coding somatic variants** |  |  |  |  |  |  |  |
| Number of sSNVs | 5426 | 4763 | 1942 | 2397 | 1381 | 1451 | 1017 |
| Number of sIndels | 199 | 202 | 85 | 94 | 68 | 52 | 25 |
| **Number of non-synonymous variants** |  |  |  |  |  |  |  |
| Missense | 3598 | 3093 | 1259 | 1566 | 898 | 964 | 650 |
| Nonsense | 223 | 188 | 78 | 82 | 42 | 58 | 37 |
| Frameshift | 196 | 192 | 80 | 93 | 67 | 51 | 24 |
| Start loss | 0 | 2 | 2 | 4 | 3 | 6 | 0 |
| Stop loss | 0 | 1 | 0 | 2 | 3 | 0 | 2 |
| Inframe deletion | 2 | 9 | 5 | 1 | 1 | 0 | 1 |

While the targeted sequencing design precluded any formal mutational signature analysis, we examined the opportunity-normalized mutational spectrum across all detected sSNVs to gain insight into the potential mutational processes. The spectrum was dominated by C>T transitions, followed by T>C transitions, consistent with spontaneous deamination and other age-related mutational processes that accumulate in post-mitotic neurons (**Supplementary Fig. 5**).^32^ C>T transitions were modestly enriched at CpG dinucleotides relative to non-CpG sites (rate ratio = 1.25, 95% CI = 1.16, 1.34; *P* = 2.42×10^-7^; two-sample Poisson rate test), with near-equal rates across all non-CpG contexts (range: 0.99–1.02), pointing to an additional contribution of 5-methylcytosine deamination. C>A, C>G and T>A transversions were comparatively low.

Across all targeted genes, 84% of neuronal nuclei (*n* = 33,542) carried at least one somatic coding variant, of which 54% carried at most two variants, while only 6% of nuclei carried more than five coding variants (median: two coding variants per neuron, range: 1–13; **Fig. 1E**). We observed substantial variability in the number of coding variants detected per sample (range: 37–6,338; **Supplementary Table 6**), which strongly correlated with the number of captured nuclei (Spearman *r* = 0.982). The somatic variant allele frequencies (VAFs), calculated as the proportion of nuclei carrying the variant among all callable nuclei at that genomic position in that sample, ranged from 0.02% to 9.09%. VAF estimates were similarly influenced by the number of captured nuclei, such that singleton variants (observed in a single neuron in a sample) appeared at inflated frequencies in samples with fewer nuclei (**Fig. 1G**). On average, 94% of variants detected in a sample were singletons (range: 68%–100%), with slightly higher singleton rates in controls (97.1%) than in patients (92.9%), reflecting differences in the number of nuclei profiled. Despite this high prevalence of singletons within individuals, >75% of somatic variants were recurrent across individuals (*n* = 14,441; **Fig. 1F**). All subsequent analyses focused on protein-coding variants and, where applicable, were adjusted for the total number of nuclei sampled per individual.

### Low-frequency singleton variants are detected throughout the *TARDBP* coding region

Our amplicons cover 88.3% of *TARDBP*’s protein-coding nucleotides (1,099/1,245) and 10 of its 16 canonical splice-site nucleotides, with full coverage of RNA Recognition Motif 1 (RRM1); partial coverage of the NTD (excluding residues 67–79), RNA Recognition Motif 2 (RRM2, excluding residues 247–263) and CTD (excluding residues 293–295 and 340–351) (**Supplementary Table 7**). We identified 1,503 somatic coding variants across 1,040 nucleotide positions, affecting 93.8% of the targeted *TARDBP* coding region, including 411 synonymous, 964 missense, 58 nonsense, 51 frameshift, 13 splice site and six start-loss variants (**Supplementary Table 8**). Variants were observed at all but eight targeted amino acids (**Fig. 2A**). The allele frequencies of somatic *TARDBP* variants ranged from 0.02% to 4.54% (median: 0.12%). Almost 80% of detected variants (1,196/1,503) were observed in more than one individual. In more than a third of the samples (30/78), all detected *TARDBP* variants were singletons (**Fig. 2C**), while the singletons, on average, accounted for 90% of the variants in the remaining samples.

**Figure 2.**
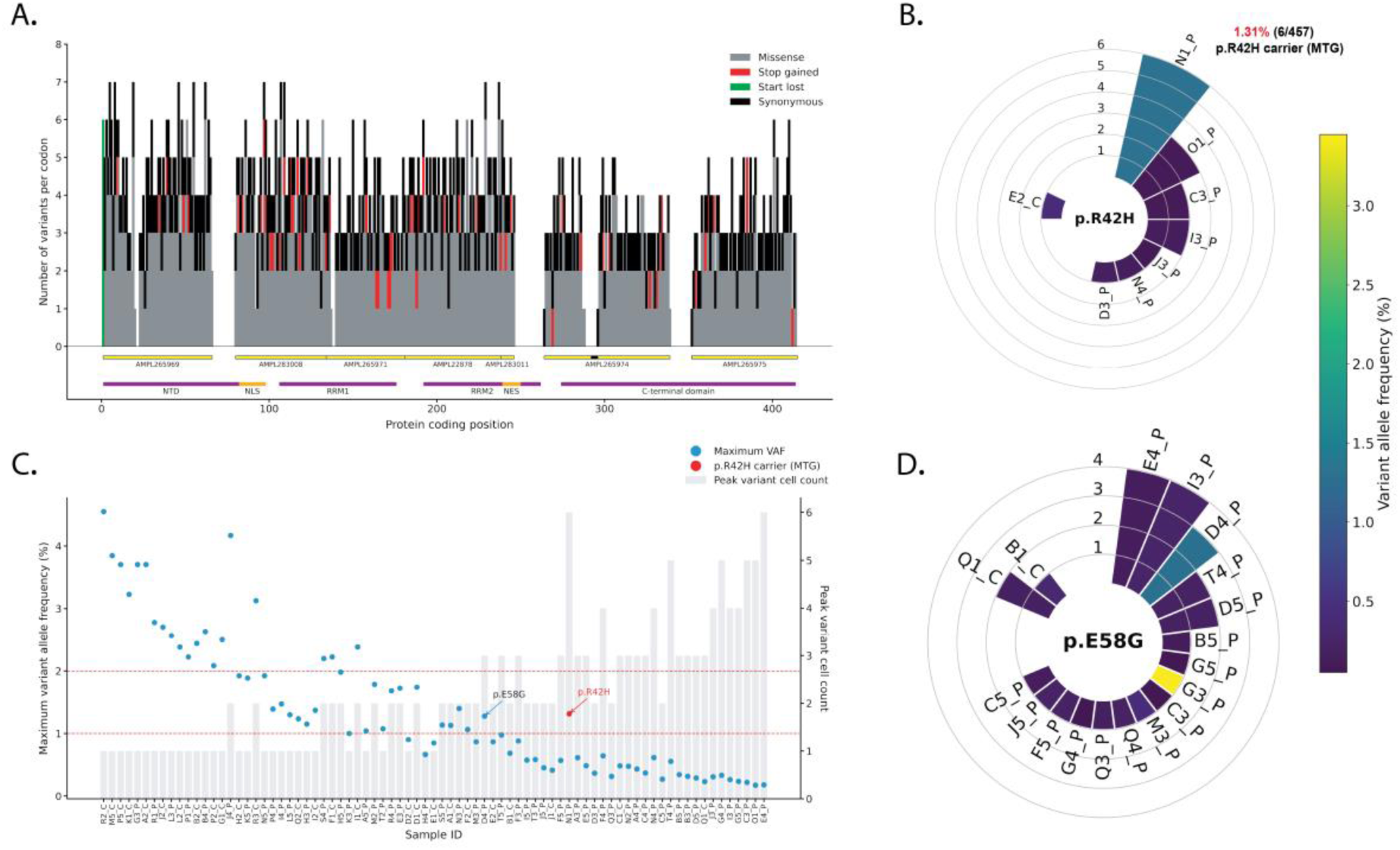
Somatic variants detected across the *TARDBP* coding sequence. **(A)** Number of somatic variants per codon plotted along the *TARDBP* protein sequence (414 amino acids). Bars are coloured by variant class (missense, stop gained, start lost and synonymous). Yellow horizontal bars indicate amplicon coverage (88% of the coding sequence; the black region indicates a 10-bp uncallable region due to insufficient read extension). Purple and orange horizontal bars denote key functional domains and motifs: N-terminal domain (NTD), RNA recognition motifs (RRM1 and RRM2), nuclear localization signal (NLS), nuclear export signal (NES) and the C-terminal glycine-rich domain (CTD). **(B)** Per-sample maximum variant allele frequency (VAF; left y-axis, blue dots) and peak variant nuclei count (maximum nuclei carrying any single variant; right y-axis, grey bars) plotted for each sample (x-axis), ordered by increasing number of total nuclei profiled. Red dashed lines indicate VAF thresholds of 1% and 2%. Samples with higher maximum VAF (>1%) predominantly carry singleton variants, whereas deeper neuronal profiling yields lower maximum VAF (<0.5%), supported by 2–6 nuclei. All samples are from the STG, except N1_P, which represents the MTG sample from the p.R42H carrier (highlighted in red). The p.R42H variant in N1_P is notably detected in six nuclei at a VAF of 1.31%, distinguishing it from other samples with comparable or more nuclei profiled. The p.E58G variant in sample D4_P (highlighted in black) is the only other *TARDBP* variant detected at a frequency ≥1% and observed in ≥3 nuclei. **(C)** and **(D)** Radial plots showing the distribution of the *TARDBP* p.R42H variant and p.E58G variant across samples, respectively. Bar lengths in both radial plots represent the number of nuclei in which the variant was detected per sample, and bar colour indicates the VAF in that sample. The *TARDBP* p.E58G variant is detected at a VAF of 1.28% in sample D4_P. Both variants are also detected at lower frequencies or in fewer nuclei in other samples, including control individuals.

We first assessed the frequency of the previously reported somatic *TARDBP* p.R42H variant in two brain tissue samples from the known carrier.^19^ In the frontal cortex, we detected the p.R42H variant at an allele frequency of 0.25% (2/795 nuclei, Test1_P from pilot experiment; **Supplementary Table 9**), close to the previously reported 0.5% in the frontal lobe. In the carrier’s MTG sample (N1_P), we observed the p.R42H variant at 1.31% (6/457 nuclei) in neurons, comparable to the previously reported 1.4% in bulk MTG tissue from the same individual. No neuronal nuclei were recovered from the STG sample of the p.R42H carrier (T1_P). The p.R42H variant was detected in the STG of six other patients at frequencies below 0.2%, in 1–3 nuclei per sample, and as a singleton in one control sample at VAF = 0.43% (**Fig. 2B**). These findings suggest that such somatic mutations may occur at very low frequencies, effectively escaping detection by bulk sequencing, which cannot discern sequencing errors from very rare somatic events.

Focusing on samples with comparable or greater numbers of sequenced neuronal nuclei than N1_P (*n* = 587), none carried a variant approaching the frequency of the p.R42H variant in N1_P (**Fig. 2C**). However, additional *TARDBP* variants with allele frequencies >1% were present in 42 samples with fewer sequenced neuronal nuclei (23 patients). To account for inflated allele frequencies in samples with few captured nuclei, we focused on variants detected in at least three nuclei within a given sample. Under these criteria, only a single variant was retained, p.E58G in patient sample D4_P, detected at a frequency of 1.28% (3/235 nuclei; **Supplementary Fig. 6**). Although p.E58G did not meet the same strict criteria in other samples, it was observed in 13 additional patients and two controls (1–4 nuclei per sample, VAF: 0.05%–0.43%; **Fig. 2D**). The p.E58G variant was also observed in sample G3_P at a VAF of 3.45% (1/29 nuclei).

Based on these findings, we hypothesized that other somatic mutations in *TARDBP* that are overrepresented or detected only in patients may contribute to disease pathogenesis. Identifying such variants could also help pinpoint regions of the gene that may be particularly vulnerable to the accumulation of somatic mutations. To identify variants overrepresented in patients, we devised selection criteria based on VAF, number of affected nuclei and functional consequence. First, we selected all non-synonymous variants detected in at least two neurons or at VAF ≥1% in a patient sample (*n* = 376). We then narrowed our selection to variants that fulfilled this criterion in at least five patients and excluded those that met the same criterion in any control sample. This yielded 14 candidate variants, all missense, with nine located in the NTD, three in the nuclear localization signal (NLS) and the remaining two in RRM2 (**Fig. 3A** and **Supplementary Table 10**). Notably, all 14 variants were predicted to be deleterious. The two most frequently observed variants were p.I18V and p.E9G. The p.I18V variant was detected in 21 patients (32 nuclei) and two controls (both singletons), whereas p.E9G was detected in 19 patients (29 nuclei) and three controls (singletons). Two variants, p.D86G and p.F211S, were detected in 15 patients (23 nuclei) and 10 patients (17 nuclei), respectively, and were not found in any control sample.

**Figure 3.**
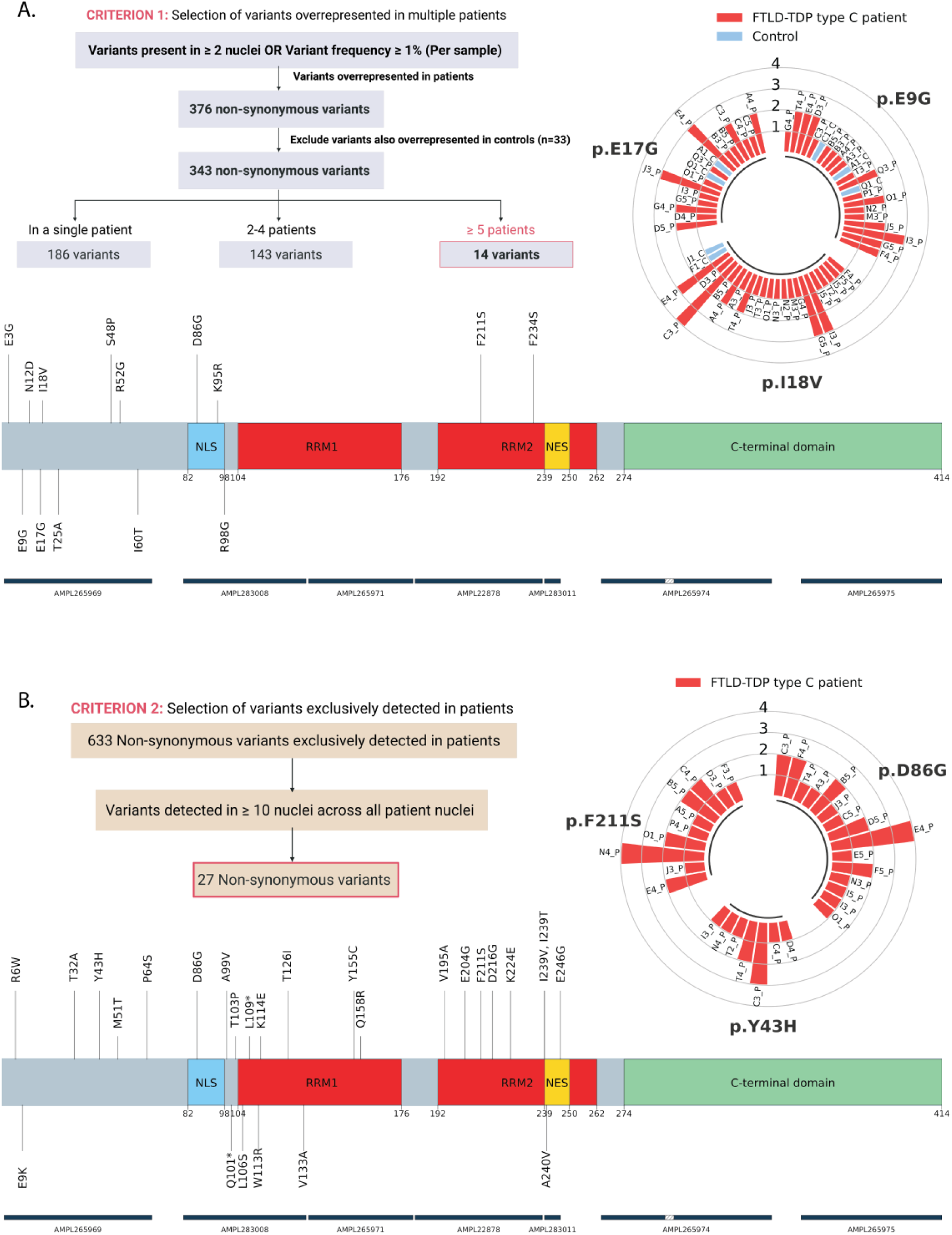
Somatic *TARDBP* variants overrepresented in or exclusive to FTLD-TDP type C patients. **(A)** Plot showing 14 non-synonymous somatic *TARDBP* variants meeting the first selection criterion: detected in ≥2 nuclei per sample or at a VAF ≥1% in at least five patient samples, with no control sample meeting the same criteria. Variants are mapped to the *TARDBP* protein sequence, with key functional domains annotated as the N-terminal domain (NTD), nuclear localization signal (NLS), RNA recognition motifs (RRM1 and RRM2), nuclear export signal (NES) and the C-terminal glycine-rich domain (CTD). Radial plots for selected variants show the distribution across samples, with bars representing the number of nuclei carrying the variant and bar colour indicating disease status. **(B)** As in (A), but applying the second selection criterion: 27 non-synonymous variants detected in ≥10 nuclei across all patient nuclei in the cohort, irrespective of per-sample thresholds. Radial plots are shown for selected variants as described above. Blue horizontal bars on both plots indicate targeted amplicon regions.

In a parallel analysis, we focused on non-synonymous variants detected in at least 10 nuclei across all patients, independent of their sample-specific frequencies. The threshold was calculated based on the 95^th^ percentile of the distribution of affected nuclei for patient-exclusive variants. This resulted in 27 variants: 25 missense (two with additional predicted splice-related consequences) and two nonsense variants, distributed across the NTD and RRMs with a notable absence of possible pathogenic somatic variants in the CTD (**Fig. 3B** and **Supplementary Table 10**). Of the missense variants, 18 were predicted to be deleterious. The most frequently observed patient-specific *TARDBP* variants included p.D86G (23 nuclei, 15 patients), p.E204G (19 nuclei, 13 patients), p.F211S (17 nuclei, 10 patients), p.E9K, p.P64S and p.D216G (each in 16 nuclei across 11 patients). Interestingly, we detected a predicted deleterious NTD variant, p.Y43H, in 10 nuclei across seven patients (1–3 nuclei per sample, VAF: 0.06%–1%). This patient-specific recurrent p.Y43H substitution is located immediately adjacent to the reported p.L41F and p.R42H variants.^19^

### Somatic mutational burden differs across TDP-43 domains

Despite the well-established pathogenic role of germline *TARDBP* CTD variants in ALS and FTD,^33–38^ no variants in the CTD met either of the two selection criteria described above. We therefore systematically assessed whether the mutational burden, defined as the proportion of neuronal nuclei harbouring at least one coding variant, differed across the TDP-43 domains. The analysis revealed significant differences in burden across domains (Bonferroni-adjusted *P* < 1×10^-16^ for all pairwise comparisons; negative binomial mixed-effects model), with higher burdens observed in the NTD (3-fold), RRM1 (2.2-fold) and RRM2 (2.9-fold) relative to the CTD (**Fig. 4A, Supplementary Table 11,** and **Supplementary Fig. 7A** and B). However, the lower burden in the CTD was not attributable to disease status, as no significant condition-specific differences at the domain level were observed (LRT: χ²(3) = 2.83, *P* = 0.42). The lower overall burden in the CTD might reflect reduced cellular tolerance to mutations in this domain, whereby neurons harbouring such mutations may be selectively lost during life and thus underrepresented at autopsy.

**Figure 4.**
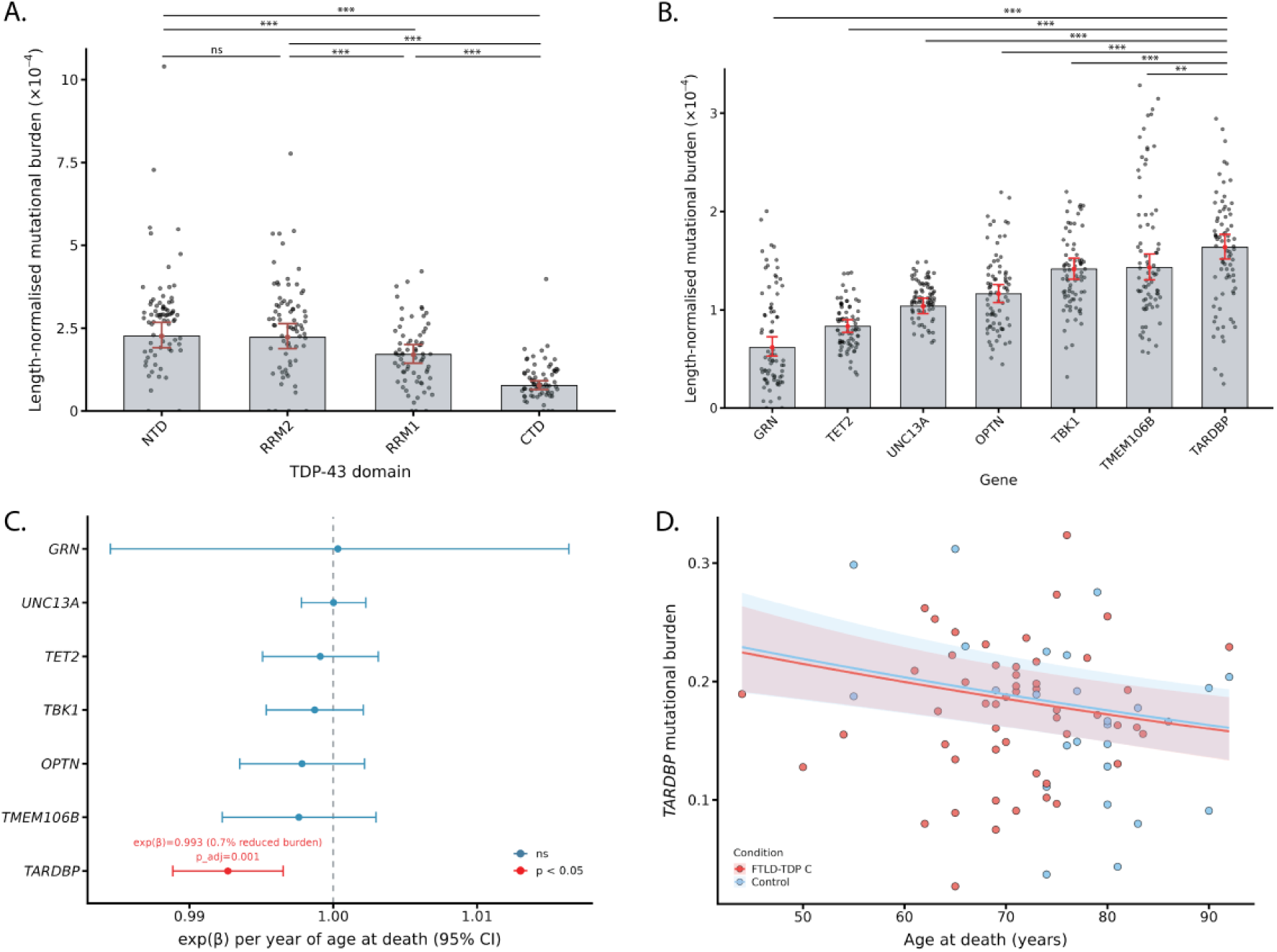
Domain- and gene-level somatic mutational burden and age-associated effects. **(A)** Per-domain and **(B)** per-gene somatic mutational burden. Each point represents the per-sample mutational burden, calculated as the number of nuclei carrying a somatic coding variant normalized for the total nuclei sampled and the targeted domain/gene length. Bars show estimated marginal means (EMMs) derived from negative binomial mixed-effects models, where mutational burden per sample was modelled as the outcome with offsets for total nuclei and gene/domain length, adjusting for disease status, age at death, sex, brain hemisphere and sequencing depth. Red points and error bars denote EMMs ± 95% confidence intervals. Pairwise comparisons are Bonferroni-corrected. *** *P* ≤ 0.001; ** *P* ≤ 0.01; * *P* ≤ 0.05; ns, not significant. **(C)** Forest plot showing the exponentiated model coefficient (β) for age at death on somatic mutational burden across targeted genes, estimated from per-gene negative binomial mixed-effects models adjusting for disease status, sex, brain hemisphere and sequencing depth, with offset for total nuclei sampled. Error bars represent 95% Wald confidence intervals. Significant genes (Bonferroni-adjusted *P* < 0.05) are highlighted in red with exp(β), percentage change in mutational burden per year of age at death and Bonferroni-adjusted *P*-value annotated. **(D)** *TARDBP* somatic mutational burden plotted against age at death, coloured by disease status. Regression lines and 95% confidence intervals are derived from a negative binomial mixed-effects model adjusting for the same covariates as in (C) and are shown at the median sequencing depth and median nuclei captured for the left brain hemisphere for male individuals.

Even with an overall underrepresentation, 300 non-synonymous variants were detected in the CTD, of which 222 were observed only in patients and nine only in controls. While 47 of these CTD variants were identified in at least two nuclei or with VAF ≥1%, each was observed in at most two patient samples, and no CTD variant was collectively detected in ≥10 patient nuclei, explaining their absence from the prioritization strategies mentioned above. The presence of these rare CTD variants prompted us to examine whether any corresponded to known ALS/FTD-associated germline mutations. For this analysis, we focused on 54 known *TARDBP* mutations retrieved from the ALSoD database,^39^ including both highly pathogenic mutations and risk variants associated with ALS/FTD. Overall, 25 of the 54 selected variants were detected at very low frequencies in our cohort: 12 were present only in patients, and 13 were present in both patients and controls (**Table 2**). Among the 12 patient-specific variants, p.N267S was the most frequent, detected in eight nuclei across six patients (1–2 nuclei per sample, VAF: 0.03%–0.25%). Other top variants included p.S393L, p.M311V and p.M337V. The latter was detected in five nuclei across three patients (1–3 nuclei per sample, VAF: 0.07%–0.18%).

**Table 2.** ALS/FTD-associated *TARDBP* variants detected somatically in the cohort. Contains 25 somatic TARDBP variants among the 54 germline ALS/FTD-associated TARDBP variants documented in the ALSoD database. For each variant, the total number of neuronal nuclei carrying the variant in patients and controls; the total number and identifiers of affected samples; and germline population frequencies from Project MinE in non-ALS individuals and gnomAD v2.1.1 in non-neurological samples are provided.

| Genomic Variant | Protein Variant | Nuclei count (Patients) | Nuclei Count (Controls) | Number of samples affected | Affected Samples | Project MinE<sup>39</sup> (Without ALS) | GnomAD v2.1.1<sup>40</sup> (Non-Neuro) |
| --- | --- | --- | --- | --- | --- | --- | --- |
| chr1:11022209:A/G | N267S | 8 | 0 | 6 | I3_P, J3_P, N2_P, O1_P, Q3_P, C3_P | 3.58E-05 | 8.17E-05 |
| chr1:11022587:C/T | S393L | 6 | 0 | 5 | G4_P, G5_P, I3_P, C3_P, F3_P |  | 1.05E-05 |
| chr1:11022340:A/G | M311V | 5 | 0 | 5 | J3_P, C3_P, C5_P, E3_P, F5_P |  |  |
| chr1:11022418:A/G | M337V | 5 | 0 | 3 | Q3_P, E4_P, F5_P |  |  |
| chr1:11022318:A/T | Q303H | 3 | 0 | 3 | G4_P, G5_P, E4_P | 7.17E-06 |  |
| chr1:11022301:G/A | G298S | 2 | 0 | 2 | N2_P, E4_P |  |  |
| chr1:11022413:G/A | G335D | 2 | 0 | 2 | N2_P, F3_P |  |  |
| chr1:11022541:A/G | N378D | 2 | 0 | 2 | Q3_P, T4_P |  |  |
| chr1:11022557:T/C | I383T | 2 | 0 | 2 | B5_P, D5_P |  |  |
| chr1:11022352:G/A | A315T | 1 | 0 | 1 | O1_P |  |  |
| chr1:11022536:G/A | G376D | 1 | 0 | 1 | E4_P |  |  |
| chr1:11022544:T/C | S379P | 1 | 0 | 1 | B3_P |  |  |
| chr1:11018836:A/G | D169G | 6 | 1 | 5 | B3_P, B5_P, E4_P, G4_P, Q1_C |  |  |
| chr1:11022484:A/G | M359V | 6 | 1 | 7 | E4_P, F3_P, G4_P, G5_P, J3_P, N5_P, C1_C |  | 4.84E-06 |
| chr1:11016874:C/T | A90V | 5 | 1 | 5 | C3_P, F3_P, M2_P, O1_P, Q1_C | 6.16E-04 | 2.18E-04 |
| chr1:11022371:C/T | A321V | 5 | 1 | 6 | A3_P, G5_P, H5_P, J3_P, M2_P, Q1_C |  |  |
| chr1:11022511:G/A | G368S | 4 | 1 | 5 | C4_P, E4_P, N4_P, T5_P, E1_C |  | 2.00E-05 |
| chr1:11022268:G/A | G287S | 3 | 1 | 4 | D5_P, N2_P, O1_P, A2_C | 3.58E-05 | 9.61E-06 |
| chr1:11022404:G/A | S332N | 3 | 1 | 4 | D4_P, E4_P, F5_P, D2_C |  |  |
| **chr1:11022553:G/A** | A382T | 3 | 1 | 4 | A4_P, B3_P, J3_P, I1_C |  | 5.28E-06 |
| **chr1:11022577:A/G** | N390D | 3 | 1 | 4 | B5_P, E4_P, G4_P, F2_C |  | 5.25E-06 |
| **chr1:11022532:A/G** | S375G | 2 | 1 | 3 | C5_P, N3_P, I2_C |  | 2.58E-05 |
| **chr1:11022556:A/G** | I383V | 2 | 1 | 2 | E4_P, Q1_C | 7.23E-06 | 1.05E-05 |
| **chr1:11022542:A/G** | N378S | 1 | 1 | 2 | C3_P, B1_C |  |  |
| **chr1:11022578:A/G** | N390S | 1 | 1 | 2 | E4_P, J1_C |  | 2.09E-05 |

Given that 13 variants were detected in both patients and controls, we examined whether these variants differed in their germline representation compared to those detected exclusively in patients. We reasoned that *TARDBP* variants, which are also found as germline variants in non-demented individuals, might be better tolerated and therefore unlikely to confer pathogenicity when observed somatically. Using non-ALS control individuals from project MinE^40^ and non-neuro samples from gnomAD v2.1.1,^41^ we found that 12 of the 25 ALS/FTD variants we had identified somatically were detected in these cohorts at extremely low minor allele frequencies (<10^-4^). Interestingly, variants exclusive to patients in our study were found less likely to be present in the population databases than those shared with control individuals (25%, 3/12 vs. 69.2%, 9/13; Odds ratio = 0.15, 95% CI = 0.026, 0.86; *P* = 0.047; Fisher’s exact test; **Table 2**), though the small number of variants warrants cautious interpretation.

### Screening somatic variants in other FTLD-TDP-associated genes

While not the primary aim of this study, we also investigated potentially pathogenic somatic variants in other FTLD-TDP-associated genes. As with the previous selection in *TARDBP* that identified p.E58G in patient D4_P, we selected variants present above 1% frequency and observed in at least three neurons (**Table 3**). *TET2* p.P1666S variant was most commonly observed in 18 patients and three controls, and additionally in six patients and three controls below the defined thresholds. However, the variant is predicted to be benign. In addition to *TARDBP* p.E58G, we identified eight variants in patients (seven missense and one frameshift): four in *UNC13A*, two in *OPTN*, one in *TET2* and one in *GRN,* each detected in a single patient sample at the selected threshold. Below the 1% VAF threshold, six of these variants were detected in both patient and control samples, predominantly as singletons, while two variants were patient-exclusive: *OPTN* p.Q277R and *UNC13A* p.Q1577R. Two variants, *UNC13A* p.R579G and *GRN* p.C247R, were identified in the same sample, F3_P, at the selected thresholds.

**Table 3.** High-confidence somatic variants detected at VAF >1% and supported by at least three nuclei. For each variant, the variant identifier, gene, sample identifier, variant allele frequency (VAF), predicted functional consequence, AlphaMissense pathogenicity score, CADD Phred score, and the number of additional samples in which the variant was detected below the selection threshold are provided. *Predicted as a high-confidence loss-of-function variant by LOFTEE.

|  | Protein Variant | Gene | Sample ID | VAF | Consequence | AlphaMissense Pathogenicity Score | CADD Phred Score | Number of affected samples (criterion not met) |
| --- | --- | --- | --- | --- | --- | --- | --- | --- |
| chr19:17627545:T/C | N1295S | *UNC13A* | P2_C | 7.89 (3/38) | Missense | 0.06 | 17.7 | 5 patients & 1 control |
| chr19:17610021:T/C | Q1577R | *UNC13A* | S4_P | 3.23 (3/93) | Missense | 0.19 | 25.0 | 3 patients |
| chr10:13116321:A/G | R203G | *OPTN* | F1_C | 3.06 (3/98) | Missense | 0.08 | 15.4 | 6 patients |
| chr10:13109265:T/A | L48Q | *OPTN* | S5_P | 2.34 (3/128) | Missense | 0.71 | 25.4 | 2 patients & 1 control |
| chr19:17621858:T/C | R1406G | *UNC13A* | M3_P | 1.68 (3/179) | Missense | 0.28 | 25.8 | 7 patients & 1 control |
| chr4:105242845:A/G | K1171R | *TET2* | F2_C | 1.47 (3/204) | Missense | 0.12 | 25.8 | 7 patients |
| chr19:17649561:T/TA | D489Vfs*210 | *UNC13A* | A3_P | 1.35 (6/443) | Frameshift* | – | – | 5 patients & 1 control |
| chr1:11013900:A/G | E58G | *TARDBP* | D4_P | 1.28 (3/235) | Missense | 0.84 | 27.3 | 14 patients & 2 controls |
| chr4:105259689:A/G | S1292G | *TET2* | M3_P | 1.28 (3/235) | Missense | 0.96 | 26.8 | 9 patients & 3 controls |
| chr10:13122435:A/G | Q277R | *OPTN* | D4_P | 1.16 (3/259) | Missense | 0.20 | 24.1 | 8 patients |
| chr19:17648512:T/C | R579G | *UNC13A* | F3_P | 1.07 (3/280) | Missense | 0.98 | 24.6 | 8 patients & 1 control |
| chr17:44351067:T/C | C247R | *GRN* | F3_P | 1.01 (3/296) | Missense | 0.96 | 29.6 | 11 patients & 2 controls |
| chr4:105275506:C/T | P1666S | *TET2* | 18 patients & 3 controls | 1.03 – 6.4 (3 – 44 nuclei) | Missense | 0.07 | 12.8 | 6 patients & 3 controls |

We then applied the same variant selection criteria used for *TARDBP* to identify patient-enriched and exclusive variants across all other FTD-associated genes (**Supplementary Table 12**). Notably, a patient-exclusive, high-confidence somatic loss-of-function *GRN* variant, p.Q249Kfs*7, was observed in 14 nuclei across eight patients (VAF: 0.1%–1.1%). This variant introduces a premature termination codon at position 255, affecting the same codon as the pathogenic p.Q249X,^42,43^ and is expected to cause progranulin haploinsufficiency.^44,45^

### *TARDBP* exhibits the highest somatic mutational burden

Finally, we compared length-normalized mutational burden across all targeted genes to assess somatic variation in *TARDBP* relative to the other genes in our panel. *TARDBP* had the highest overall somatic mutational burden (Bonferroni-adjusted *P* < 0.05 for all pairwise comparisons; negative binomial mixed-effects model), followed by *TMEM106B*, *TBK1*, *OPTN*, *UNC13A*, *TET2* and *GRN*, with *TARDBP*’s burden ranging from 1.14-fold to 2.64-fold higher than the other genes (**Fig. 4B, Supplementary Table 13** and **Supplementary Fig. 7C** and **D**). We then assessed gene-specific covariate effects on mutational burden and found no significant differences between patients and controls (**Supplementary Fig. 7E**) or between sexes across any targeted gene. However, higher age at death was associated with a small but significantly lower mutational burden only in *TARDBP,* corresponding to ∼7.1% fewer variant-carrying neurons per decade of age at death (*β* = −0.00735, exp(*β*) = 0.993 per year, 95% CI = 0.989, 0.997; Bonferroni-adjusted *P* = 0.0014; negative binomial mixed-effects model) (**Fig. 4C, Supplementary Table 14**). This association was also observed in analyses stratified by disease status for *TARDBP* (**Fig. 4D**), with no statistical evidence of an interaction between age at death and disease status (LRT: χ²(1) = 0.99, *P* = 0.32).

## Discussion

In this study, we investigated the potential contribution of somatic mutations in *TARDBP* and other FTLD-TDP-associated genes to sporadic FTLD-TDP type C pathology using single-nucleus amplicon sequencing. A major challenge in studying somatic mutations in neurodegeneration is the progressive neuronal loss prior to autopsy, which reduces the number of mutation-harbouring cells and thus lowers VAFs. In this context, a key strength of our study is the scale of neuronal profiling achieved. With 39,800 nuclei sequenced across 78 FTLD-TDP type C patients and control individuals, our dataset considerably exceeds the number of neurons previously assayed in scWGS studies.^21–25,32,46–48^ The combination of single-nucleus resolution and sampling depth enabled the identification of ultra-low-frequency variants, including those present in only a single neuron, which would be difficult to distinguish from errors with bulk sequencing approaches.

We leveraged the Mission Bio Tapestri platform and demonstrated its utility in a neurodegenerative brain disorder. The platform’s design for high-throughput, targeted single-cell genotyping in haematological malignancies and solid tumours meant that analytical pipelines, variant-calling thresholds and quality-control frameworks had been optimized for tumour cell populations, which differ substantially from post-mitotic neurons in cell size, nuclear integrity, DNA quality and expected VAFs (e.g., clonal cell proliferation vs. selective neuronal degradation). By combining NeuN+ nuclei sorting with single-nucleus amplicon sequencing in a pooled-sample design, we adapted this technology to post-mortem brain tissue and developed a robust analytical framework for detecting low-frequency neuronal somatic variants. Our amplicon design achieved high per-nucleus coverage, thereby substantially reducing stochastic noise and enabling characterization of somatic variation at a resolution not previously achieved in FTLD-TDP.

Our data revealed a strikingly high number of low-abundant neuronal somatic variants in all analysed genes, including *TARDBP*, suggesting that previous studies relying on bulk sequencing or small-scale single-cell approaches significantly underestimated their abundance. On average, we observed two sSNVs per neuron across a targeted panel of seven genes. Although our panel is not a representative genomic sample, if extrapolated, this would substantially exceed prior genome-wide estimates at a similar age at death (∼1,000–9,000 sSNVs/neuron^21,23,49^). This observed per-neuron mutational burden can likely be attributed to two factors: (i) our gene-level focus on expressed loci, which are expected to show elevated mutational burden^32^; and (ii) our large-scale profiling of neurons, which provides greater sensitivity for detecting somatic variants.

Importantly, we confirmed the presence of the somatic *TARDBP* p.R42H variant, previously implicated in FTLD-TDP type C, in two brain tissues from the known carrier at frequencies closely matching those observed with deep bulk sequencing. Beyond p.R42H, *TARDBP* p.E58G was identified as another variant in an FTLD-TDP type C patient, with an allele frequency exceeding 1% and present in at least three neurons (VAF = 1.28%). Both variants were also detected in additional patient and control samples at frequencies below 0.5%, suggesting that N-terminal somatic variants may constitute part of the normal landscape of somatic variation in *TARDBP*, which may reach a critical threshold in some individuals and potentially contribute to disease.

An important limitation, which only became apparent during data analysis, complicated the interpretation of disease-associated variants in our study. We noticed that our pooled libraries unexpectedly recovered proportionally fewer nuclei from controls than patients. Since such an imbalance was not observed in the unsorted pilot experiment (**Supplementary material, ‘Pilot experiment’ section**), it was likely introduced during the neuronal enrichment steps, possibly reflecting disease-associated differences that favour neuronal capture of patient nuclei over control nuclei, though the precise mechanism remains unclear. As a result, VAFs were inherently inflated in the more sparsely sampled control population, hindering our ability to use them alone as a discriminator of disease relevance. Consequently, we devised variant selection strategies focused on variants enriched or found exclusively in patient neurons, reasoning that such variants may better represent disease-relevant candidates.

These filtering strategies prioritized several *TARDBP* variants. Multiple variants affected the NTD, including the patient-specific *TARDBP* p.Y43H. Combined with the previous discoveries of p.L41F and p.R42H, the recurrence of somatic mutations in this region may constitute a mutational hotspot that may affect NTD’s function in TDP-43 folding and self-oligomerization.^50–52^ We also identified variants in the NLS and RRMs, with *TARDBP* variants p.D86G and p.F211S prioritized under both variant selection strategies. Neither of these variants was previously associated with ALS/FTD. The *TARDBP* p.D86G maps to the NLS and may partially disrupt nuclear localization, while p.F211S affects a highly conserved RRM2 residue. In addition to p.D86G, we found two other patient-enriched *TARDBP* variants in the NLS: p.K95R and p.R98G. In the germline, p.A90V remains the only ALS-associated *TARDBP* variant within the NLS shown to cause TDP-43 mislocalization *in vitro*, and it has been proposed as a risk factor for ALS/FTD rather than a fully penetrant pathogenic mutation.^53,54^ The somatic NLS variants identified here may similarly confer disease risk in affected individuals.

A striking observation that emerged from our analyses was the relatively low mutational burden in the CTD compared with other TDP-43 domains. This appears counterintuitive given that most germline pathogenic ALS/FTD mutations cluster within the C-terminal region. We suggest this may reflect survivor bias, whereby neurons carrying highly deleterious CTD mutations are progressively lost during life and therefore become underrepresented at autopsy. In fact, upon focused screening for known ALS/FTD-associated variants, we did detect rare occurrences of such CTD variants in patient nuclei, including well-established pathogenic germline mutations such as p.N267S and p.M337V. These variants were detected at frequencies below the thresholds for classification as either patient-enriched or patient-exclusive variants. This may also explain why previous analysis of somatic variants in FTLD-TDP type C using bulk sequencing failed to identify any *TARDBP* mutations known to cause disease in the germline.^19^ If survivor bias accounts for the low frequencies at which CTD variants are observed in post-mortem tissues, we expect more neurons with these mutations to be present earlier in the disease course, where they may contribute to, or potentially drive, disease pathogenesis. When affecting sufficient numbers of neurons, these pathogenic mutations may initiate or confer disease risk by promoting TDP-43 mislocalization and aggregation, a process that could be further amplified through prion-like propagation to neighbouring neurons.^55–61^ This hypothesis is supported by a recent report of a highly focal somatic *TARDBP* p.L248F variant (VAF = 0.75%) in a sporadic ALS/FTD patient, where the anatomical site of variant discovery was associated with an elevated phospho-TDP-43 burden.^17^

We also examined other FTLD-TDP-associated genes, primarily to contextualize the *TARDBP* findings, rather than to detail their somatic contribution to FTLD-TDP type C. Germline mutations in these other genes have been implicated as causal or risk factors for FTLD-TDP subtypes A and B,^44,45,62–67^ but not FTLD-TDP type C. While a few likely deleterious somatic variants with VAFs exceeding 1% and present in at least three nuclei were identified in *UNC13A, OPTN* and *GRN*, these specific variants had no prior germline disease association, and their functional relevance remains unknown. Importantly, *TARDBP* was found to be the only gene with a lower mutational burden in individuals with a higher age at death. Since directionality is unknown, this may reflect that individuals dying at a younger age carry a higher *TARDBP* mutational burden. However, it may also reflect progressive loss of neurons carrying pathogenic *TARDBP* mutations over time, accounting for their reduced prevalence in older individuals. This is further corroborated by *TARDBP*’s high germline constraint (missense o/e ratio = 0.39 and pLI = 1),^41^ indicating that coding variation in this gene is poorly tolerated, an observation that may extend to somatically arising variants. While a genome-wide increase in somatic mutational burden with age, as observed in other neurodegenerative diseases,^21–23,25,32^ remains plausible, our data demonstrate that gene-specific changes in burden may go undetected by scWGS at limited sequencing depth, underscoring the advantage of our approach.

Furthermore, *TARDBP* exhibited the highest overall mutational burden among all the genes examined. Highly expressed genes accumulate more neuronal somatic mutations arising primarily from transcription-associated mutagenesis.^32,46,47,49^ Although expression profiles from the STG are unavailable, single-cell transcriptomics data from the MTG show that all targeted genes are highly expressed in neurons: *UNC13A, OPTN, TMEM106B, TET2* and *TBK1* rank within the top 10%, followed by *TARDBP* in the top 20% and *GRN* in the top 40%.^68,69^ Therefore, the elevated susceptibility to somatic *TARDBP* variation cannot be attributed to transcriptional activity alone, suggesting contributions from additional factors such as local sequence context, DNA repair efficiency, or chromatin accessibility. Given the potential influence of survivor bias, disease-specific effects cannot yet be excluded and will likely require substantially larger sequencing efforts, both in terms of the number of individuals and neuronal nuclei profiled per individual.

Based on the widespread occurrence of low-frequency somatic mutations across all targeted genes, it remains possible that a subset of somatic variants identified in this study arise as a molecular consequence of FTLD-TDP pathology, as loss of nuclear TDP-43 has been shown to impair DNA repair and promote genomic instability.^70–72^ Disease pathology is a known mutagenic driver in many neurodegenerative diseases. Oxidative damage has been linked to an increased genome-wide burden of neuronal somatic mutations in Alzheimer’s disease,^21^ and more recently in *C9orf72*-associated ALS and FTD,^23^ while neuroinflammation has been similarly implicated in multiple sclerosis.^22^ However, exploring this possibility would require either single-nucleus sequencing of broader gene panels including multiple dementia-associated genes or genome-wide profiling.

Future efforts could be directed toward screening somatic *TARDBP* mutations across multiple tissue types from the same individuals, which could help establish whether certain variants arise during early development or are restricted to specific brain regions, providing insight into their developmental versus post-mitotic origin. In view of the recent discovery of heteromeric annexin A11-TDP-43 amyloid fibrils in FTLD-TDP type C,^10,11^ a finding not known at the time our panel was designed, future studies could also explore somatic *ANXA11* mutations. Furthermore, systematically establishing the functional consequences of somatic variants across all TDP-43 domains, extending previous efforts focused on its CTD,^73^ will be crucial for interpreting their pathogenic relevance. In this context, approaches such as massively parallel reporter assays or saturation genome editing could be employed to investigate the effect of all possible *TARDBP* mutations on the splicing regulation of its key downstream targets, such as *UNC13A* and *STMN2.*^74–77^

In conclusion, we show that targeted single-nucleus sequencing of a large number of neurons enables detection of ultra-low-frequency somatic variants, including those present in only a single neuron per individual, providing the first evidence of gene-specific somatic mutations at single-neuron resolution in sporadic FTLD-TDP. The extent and range of the observed variants, spanning nearly all codons of the targeted genes, are striking and highlight the extensive accumulation of somatic mutations in post-mitotic neurons, not only in diseased individuals but also in neuropathological controls. We identify multiple candidate disease-associated somatic mutations, including *TARDBP* variants with established roles in ALS and FTD, variants that would likely go undetected by bulk or single-cell approaches lacking sufficient sequencing depth. Together, these findings establish targeted single-nucleus sequencing as a powerful approach for detecting ultra-low-frequency somatic variants at scale and provide a framework for future studies aimed at defining the contribution of somatic mutations to FTLD-TDP pathogenesis.

## Data and code availability

A detailed list of all somatic *TARDBP* variants, along with sample-level frequencies and counts of affected nuclei per sample, is provided in **Supplementary Table 8**. Custom scripts and code to reproduce the findings in the study are publicly available in a GitHub repository https://github.com/vanshikabidhan/somatic-ftd under the MIT license.

## Acknowledgements

We thank the VIB Nucleomics Core for support with FANS-based sorting of NeuN+ nuclei and the Biobank Antwerp, Belgium (ID: BE 7103003100).

## Author contributions

R.R. and W.D.C. conceptualized and designed the study. V.B. performed all formal data analyses. T.S., S.W., and C.T.V. optimized the protocol. S.W. performed the experiments. J.v.R, M.O.M., L.D.K., S.A.S., I.B., A.K., C.T., C.P., J.S., D.R.T., R.V., M.V., A.T.N., R.R.R., J.K., O.L.L., C.L.W., B.F.B., N.R.G., K.A.J., R.C.P., N.B.G., M.E.M., D.W.D., J.C.v.S., and H.S. collected and provided tissue samples and clinicopathological data. S.W. and M.V.d.B. acquired and processed the samples. F.K. and K.S. advised on statistical methodology. V.B., W.D.C., and R.R. drafted the manuscript. W.D.C. and R.R. acquired funding and supervised the work. All authors contributed to reviewing and revising the manuscript and approved the final version.

## Supplementary material

**Supplementary Information:** Supplementary Text and Figures 1-7.

**Supplementary Tables:** Supplementary Tables 1-14.

