## Supplementary Material for "Targeted single-nucleus sequencing of 39,800 neurons reveals extensive low-frequency somatic variants"

### Supplementary text

#### Pilot experiment

Prior to the main experiment, a pilot run was performed by pooling approximately 30 mg of frontal cortex (FCX) tissue from two FTLD-TDP type C patients (Test1_P, Test4_P) and three non-demented controls (Test2_C, Test3_C, Test5_C; **Supplementary Table 9**). Nuclei were sequenced without prior MARS-based debris removal or NeuN+ sorting. Test1_P in this experiment represents the FCX tissue sample from the same patient with the previously confirmed *TARDBP* p.R42H somatic mutation, and the variant was detected at a VAF of 0.25% (2/795) in this FCX sample.

Comparing this pilot run with the main experiment presented in the manuscript yields a few key conclusions. An average of 771 nuclei per sample were recovered from this pilot run (range: 596–992; **Supplementary Table 9**). Notably, nuclei counts from control samples (823, 655 and 596) were comparable to nuclei counts from patient samples in this unsorted run (992 and 788). This contrasts with the NeuN+-sorted nuclei from controls in our main experiment, which consistently yielded fewer nuclei than those from patients. This finding, though based on a single experiment, points to the NeuN+ sorting stage as a potential source of the reduced neuronal yield in controls as compared to patients when using the NeuN+-sorted fractions. Moreover, the pilot run yielded an average of 4,039 variants per sample, with 91% occurring as singletons, indicative of widespread somatic mosaicism, which we attributed in part to possible contributions from non-neuronal cell types. We thus reasoned that restricting variant detection to neurons might enrich for likely pathogenic variants at higher VAFs, which would otherwise be diluted by non-neuronal cell-type contributions in the unsorted nuclei population. For this reason, we implemented NeuN+ sorting in the main experiment.

Importantly, Test4_P and Test5_C were included in the NeuN+-sorted run as samples Q4_P and Q5_C, respectively, to allow comparison between the pilot run and the main experiment. In particular, for these two FCX tissue samples common to both the unsorted and NeuN+-sorted experiments, we compared the allele frequencies of variants shared between the samples. Only ~16% of variants detected in NeuN+-sorted nuclei were also called in unsorted samples, consistent with dilution of neuronal-specific variants (especially singletons) in the unsorted run. These shared variants showed higher VAFs in neuronal nuclei than in unsorted nuclei in both patient (88%, 352/402 variants) and control samples (98%, 77/79 variants), consistent with neuronal enrichment.

#### Germline variants detected in sporadic FTLD-TDP type C patients

All patients in this cohort were neuropathologically diagnosed with FTLD-TDP type C. In our analyses, we identified no known pathogenic germline variants in any gene in our panel. We detected 14 unique germline mutations in FTLD-TDP-associated genes (excluding *TET2*) in six controls and 10 patients. However, based on their population frequency, *in silico* predicted deleteriousness and known disease biology, none were expected to be pathogenic (**Supplementary Table 5**). Of the 10 germline variants identified across 10 patients, four were intronic, three missense and three synonymous. Eight were present in gnomAD (exomes and genomes, AF: 0.0003–0.02), consistent with rare-variant status. Only four variants (*UNC13A* p.V740= and p.L1184=; *TBK1* p.H322Y and p.F45=) were catalogued in ClinVar, all classified as benign. Additionally, none of the three missense variants reported in patients were predicted to be deleterious (based on both AlphaMissense pathogenicity and CADD Phred scores).

### Supplementary materials and methods

#### Cohort demographics

The 93 STG samples (**Supplementary Table 1**) included 41 female samples (21 patients, 20 controls) and 52 male samples (36 patients, 16 controls). The median age at death was 71 years (range: 44–92 years) in patients and 77 years (range: 54–92 years) in controls. The median age at disease onset in patients was 61 years (range: 42–78 years), and the median disease duration was 9.6 years (range: 2–23.6 years). In total, right-brain hemispheres were collected from 61 individuals (41 patients, 20 controls) and left-brain hemispheres from 32 individuals (16 patients, 16 controls).

#### Amplicon panel design

We designed a targeted panel of 193 amplicons (**Supplementary Table 2**): 128 covering the exonic regions of *TARDBP*, *UNC13A*, *TBK1, OPTN*, *GRN*, *TMEM106B* and *TET2*; 57 targeting a set of highly frequent germline SNPs in intergenic regions to allow demultiplexing of the pooled samples; and eight amplicons targeting chromosome Y loci for the detection of mosaic loss of chromosome Y (LOY). These germline SNPs were selected from gnomAD[1] v2.1.1 genomes by retaining variants with non-Finnish European allele frequencies between 0.4 and 0.6. Variants located in repeat-masked or segmental duplication regions, as well as indels and non-PASS filter entries, were excluded. 50 SNPs were randomly sampled from the 545,239 high-confidence sites. Of these, 43 were incorporated into the final panel due to amplicon design constraints (**Supplementary Table 3**).

For *TARDBP* specifically, exons 3 and 4 were fully covered, while exons 2, 5 and 6 were partially covered (excluding 41, 2 and 103 nucleotides, respectively). At the protein domain level, this corresponds to no observations of amino acid residues 67–79 within the NTD, full coverage of the RRM1 domain, exclusion of residues 247–263 within the RRM2 and exclusion of residues 340–351 in the CTD (**Supplementary Table 7**). The panel was originally designed against the hg19 reference genome, and the coordinates were lifted over to GRCh38 prior to downstream analyses using UCSC LiftOver.[2]

### Nuclei isolation

Brain tissue samples, weighing approximately 30 mg each, were pooled in groups of five, except for sample O1_P that contained a single sample weighing 147 mg. All samples were kept on dry ice during processing. Nuclei were isolated using the Mission Bio nuclei extraction protocol with minor modifications. Briefly, tissues were mechanically disrupted and incubated in lysis buffer (0.03 mg/mL) for 2–3 minutes, then further minced to obtain fine fragments. Samples were thawed and gently agitated at 20 rpm for 10 minutes at room temperature. After lysis, a stop solution was added, and the mixture was incubated for 1 minute at room temperature. The resulting nuclear suspension was filtered through a 40 µm cell strainer and centrifuged at 500 × g for five minutes at room temperature. Pellets were resuspended in 1 mL of spermine buffer containing 0.1% BSA. The suspension was then filtered through a 20 µm strainer and washed with 2 mL of spermine buffer containing BSA. The final nuclei suspension was kept on ice until further processing.

### Nuclei enrichment and neuronal sorting

To reduce debris and dead cells, nuclei suspensions underwent acoustic cleanup using the MARS system (Applied Cells). The acoustic frequency was optimized for each module (13.5–16 Hz). Suspensions were loaded onto the device and processed using a single workflow comprising a 5-minute wash cycle followed by a 10-minute cleanup. Nuclei were counted with the LUNA-FL automated cell counter. Following cleanup, nuclei were incubated with an anti-NeuN antibody (1:125; Milli-Mark® Anti-NeuN-PE, clone A60) for 30 minutes on ice. Nuclei were sorted via fluorescence-activated nuclei sorting (FANS) on a BD FACSAria Fusion cell sorter, isolating approximately 350,000 NeuN+ nuclei per pool. Sorted nuclei were centrifuged at 450 × g, and the resulting pellets were resuspended, combined and diluted with Mission Bio cell buffer. Nuclei were stained with Acridine Orange/Propidium Iodide (1:10) and counted using the LUNA-FL cell counter. Finally, suspensions were diluted to 3,000–4,000 nuclei/µL using cell buffer.

### Single-nucleus DNA library preparation and sequencing

Libraries for single-nucleus DNA sequencing were prepared using Mission Bio's Tapestri v2 chemistry. Nuclei were first encapsulated into droplets using the Tapestri microfluidics DNA cartridge. Within each droplet, nuclei were lysed and digested, releasing genomic DNA.

Barcoding was performed by introducing barcoding beads and a barcoding mix containing gene-specific forward primers. Barcoded nuclei underwent targeted PCR amplification with reverse primers across 193 targets from our custom-designed panel. Following amplification, intact emulsions were broken using an extraction agent, DNA-binding proteins were digested, and PCR products were purified with AMPure XP beads. Indexing was performed via a second round of PCR using i5/i7 index primers, followed by a second AMPure XP purification. Final libraries were stored at −20°C until sequencing. Library quality was assessed using Agilent Bioanalyzer. In cases where primer dimer contamination was detected, libraries were subjected to an additional cleanup using a 0.72× AMPure XP bead ratio to selectively retain larger fragments. Cleaned-up PCR products were quantified using Qubit, and all libraries were pooled equimolarly to 10 nM and sequenced on the Element Biosciences AVITI platform. Sequencing was performed targeting 1,000 nuclei per sample (5,000 nuclei per pool), yielding a mean capture of 2,360 nuclei per pool (**Supplementary Table 4**).

Raw sequencing data was processed using Mission Bio's Tapestri analysis pipeline (v2.0.2). We modified a single parameter to disable GATK downsampling and reduce false positives by setting `max-reads-per-alignment-start` to 100,000. Briefly, adapter sequences were trimmed from the reads, barcodes were extracted and error-corrected, and the reads were aligned to the human reference genome (hg38). Reads lacking an insert or mapping to an off-target site were filtered out. Barcodes with a data completeness rate of ≥80% were classified as cells. Genotyping was performed using GATK HaplotypeCaller in GVCF mode, followed by joint genotyping with GenotypeGVCFs, which retained both variant and non-variant loci in the output. The resulting output includes genotype (GT) information for all genomic variants across all nuclei, along with associated metrics such as allele frequency (AF), read depth (DP), and genotype quality (GQ). Genotype calls are encoded as follows: homozygous reference (GT = 0), heterozygous (GT = 1), homozygous alternate (GT = 2) and missing call (GT = 3). Following initial processing with the Tapestri pipeline, data were analysed using Mission Bio's mosaic pipeline (version 3.12).

### Sample demultiplexing

#### Generating reference SNP profiles through germline genotyping of demultiplexing variants

To demultiplex the pooled samples, we used the frequent germline variants included in the design and generated reference SNP profiles for all included individuals. For four samples, genotypes were obtained from in-house whole-genome sequencing data: three from short-read (L5_P, M2_P and T3_P) and one from long-read (T5_P, sequenced on Oxford Nanopore Technologies PromethION and variants called using Clair[3]), all of which were brain-derived. For the remaining samples, targeted genotyping of 34 germline SNPs (out of the 43 demultiplexing SNPs included in the panel design) was performed using multiplex amplicon sequencing on an Oxford Nanopore Technologies Flongle flow cell, followed by minimap2[4] alignment and Clair3 v0.4.0[3] variant calling. After excluding variants with insufficient coverage (>5 samples) or genotypes discordant with overlapping short-read WGS data ($n$ = 8), 26 high-confidence germline SNPs were used to generate a reference genotype profile.

#### Identification of true sample-representative cells

In each pool, the 26 high-confidence demultiplexing germline SNPs were filtered to retain those genotyped in at least 85% of nuclei. Genotypes of individual nuclei were then compared against the reference profiles of all samples within the pool. Nuclei with exact genotype matches were directly assigned to the corresponding sample. These nuclei served as true positives for downstream cluster assignment and were also used in subsequent doublet-detection analyses. To avoid ambiguity, they are referred to as "*sample representative*" nuclei. Samples with fewer than five representative nuclei ($n$ = 15) were excluded from downstream analyses as they could not be confidently deconvoluted (**Supplementary Table 6**).

#### Principal component reduction and clustering analysis

Starting with the complete set of 43 demultiplexing variants retained to maximize information for clustering, low-confidence genotypes (DP <10 and GQ <30) were set to missing, and SNPs genotyped in <80% of nuclei per pool were excluded. The resulting filtered variant set was used for principal component analysis. Nuclei with missing genotypes for more than 10 of the selected SNPs were marked as unreliable and excluded from further analysis due to insufficient data for accurate assignment ($n$ = 394, median: 7.5 per pool; **Supplementary Table 4**). Before clustering, missing genotypes were imputed by assigning each variant the mean genotype across all nuclei within the respective pool. Genotypes were discretized as follows: a mean value $\leq 0.5$ was assigned as homozygous reference (GT = 0), a value >0.5 and <1.5 as heterozygous (GT = 1) and a value ≥1.5 as homozygous alternate (GT = 2). The resulting SNP-genotype matrix was subjected to principal component reduction, retaining components that cumulatively explained ≥80% of the variance, with a minimum of five components enforced. Clustering was performed using k-means, with $k$ set to $n + 1$, where $n$ is the expected number of donor clusters, plus an additional cluster for potential mixed-origin nuclei (e.g., doublets or low-confidence nuclei). A default of $k = 6$ was applied across most pools, with pool-specific adjustments made where necessary (**Supplementary Table 4**). The sample identity for each cluster was determined by the predominant composition of sample-representative nuclei within that cluster. Cluster-to-sample concordance was assessed using Hamming distances between consensus cluster genotypes and reference profiles across the 26 filtered SNPs (**Supplementary Table 6**). Upon inspection, each discordant position reflected either a missing genotype in the reference profile or a dropout in the sequencing data. All samples scored below 5, except for T5_P ($d = 8$), where all discordant positions reflected missing genotypes in the long-read genome sequencing-derived reference profile. Nuclei clusters that could not be confidently assigned to any donor sample were labelled as unassigned.

#### Doublet removal

A doublet removal strategy was then employed to remove potential heterotypic doublets (droplets containing nuclei from two different samples). Approximately 10% of artificial doublet nuclei were introduced by combining representative nuclei from distinct donor samples identified in the initial genotype-matching step. These artificial doublets were subjected to k-means clustering, with $k$ set to $2n$, where $n$ is the number of donor samples in the pool, to accommodate all potential doublet combinations. Following clustering, the Euclidean distance of each nucleus to the centroid of each artificial cluster was calculated. Nuclei closer to an artificial doublet cluster than to the centroid of their assigned cluster were flagged as doublets. This approach removed both unassigned and incorrectly assigned nuclei. Among sample-representative nuclei, doublets were largely absent or below 1% across most samples, with five samples exhibiting rates between 2%–8% (**Supplementary Table 4**).

#### Variant filtering

Following removal of doublets and low-quality nuclei, per-genotype filtering was applied (i.e., per variant, per nucleus), setting genotypes to missing where DP <10, GQ <30 or VAF thresholds were violated (i.e., if a genotype labelled as homozygous reference has an alt-VAF >10%, a homozygous alternative genotype has VAF <85% or a heterozygous variant has a VAF <30%). Variants genotyped in fewer than 50% of nuclei within a sample were converted to missing, and a union of variants detected across all samples was subsequently taken. Variants were then excluded if they mapped to poor-performing amplicons (mean DP <10 in ≥20% of nuclei) or to sex chromosomes. Nuclei with a low average coverage (mean DP <10 across ≥20% of remaining amplicons) were subsequently removed (*n* = 236), along with variants harbouring spanning deletion alleles. Remaining variants were annotated using the Ensembl Variant Effect Predictor (VEP, release 115),[5] integrating Combined Annotation-Dependent Depletion scores (CADD v1.7),[6] AlphaMissense pathogenicity scores,[7] gnomAD population frequencies[1] and SpliceAI predictions.[8] Loss-of-function variants were annotated as high-confidence using Loss-Of-Function Transcript Effect Estimator (LOFTEE).[1] Variants were considered predicted deleterious if they had an AlphaMissense pathogenicity score >0.564 and/or a CADD Phred score >20. Variants were subsequently filtered to retain only those mapping to the target genes of interest. Variants overlapping repetitive genomic elements were excluded based on RepeatMasker annotations[9] retrieved from the UCSC Genome Browser (hg38).[2]

Per-variant callability in a sample was calculated as the proportion of nuclei genotyped at a given locus. Variants with callability below a 70% threshold were converted to missing in that sample. VAF in a sample was then calculated as the proportion of callable nuclei carrying the variant, including both heterozygous and homozygous genotype calls. Variants with a global minor allele frequency >1% in gnomAD or 1000 Genomes Project Phase 3 data were excluded to remove common germline polymorphisms. Variants detected in ≥65% of the nuclei in a sample were flagged as putative germline mutations. For samples with available genome sequencing data, germline status was confirmed orthogonally (**Supplementary Table 5**). Flagged germline variants were excluded, with their genotypes set to missing in the relevant samples. Insertions exceeding 5 bp in length were also removed.

For each heterozygous genotype call with an allele frequency below 50%, a one-sided binomial test was applied to assess whether the observed alternate allele count was significantly lower than expected under a true heterozygous state ($p_0 = 0.5$). To control for multiple testing, the Benjamini–Hochberg (BH) false discovery rate correction was applied per nucleus across all heterozygous calls within that nucleus. Heterozygous genotypes with BH-corrected $P$-values $< 0.05$ were converted to homozygous reference, reflecting allelic imbalance inconsistent with a genuine heterozygous call. Finally, variants detected within a 5-bp window in the same nucleus were set to missing to minimize the influence of primer-misalignment artefacts (**Supplementary Figs. 2 and 3**). Variants of interest were visualized using the Integrative Genomics Viewer.[10]

#### Selection of ALS/FTD-associated variants from ALSoD

A set of 69 ALS/FTD-associated *TARDBP* variants was retrieved from the ALS Online Database (ALSoD).[11] Non-synonymous SNVs were retained, yielding 58 candidates. Four variants not captured by our sequencing panel were subsequently excluded, leaving a final set of 54 variants for analysis, comprising 53 missense and one nonsense variant.

#### Estimating the somatic loss of the chromosome Y (LOY) in males

Eight amplicons targeting regions on both arms of chromosome Y were included in the amplicon panel. Amplicons were first filtered using a mean read depth threshold (DP <10 across all samples), resulting in the exclusion of one amplicon (AMPL293807). Nuclei were classified as exhibiting LOY if the selected Y-linked amplicons had zero read coverage, with a tolerance of up to three non-specific reads in a subset of amplicons. To ensure that LOY classifications reflected actual loss of signal, coverage across Y-linked amplicons in nuclei classified as LOY was compared to the mean read depth of both non-Y amplicons within the same nuclei and to Y-linked amplicons in nuclei without LOY.

#### Characterizing mutational profiles

All sSNVs detected across target genes were classified into the 96 pyrimidine-centred trinucleotide substitution contexts following COSMIC conventions. Observed mutation counts were aggregated by context. Substitution rates were normalized by the number of callable trinucleotide opportunities within the BED-defined amplicon intervals, computed by scanning all trinucleotides overlapping the targeted regions and normalizing to the pyrimidine-centred frame. Final substitution rates were expressed as mutations per million trinucleotide opportunities. C>T mutations at CpG dinucleotides were identified as contexts where the right-flanking base was G, and their mean rate was compared with that of non-CpG C>T contexts. The enrichment ratio and 95% confidence interval were estimated using a log-normal approximation of the Poisson rate ratio, and statistical significance was assessed via a two-sample Poisson rate test.

#### Statistical analyses

Negative binomial mixed-effects models were fitted to compare mutational burden separately across genes and TDP-43 domains. Both models included disease condition, age at death (mean-centred, mean = 72.53 years), biological sex, gene/domain identity, brain hemisphere and the sample-level median sequencing depth per gene/domain as fixed-effect covariates. Identifiers for the sample, sequencing pool and recruitment site were included as random effects to account for repeated measures within individuals and technical variability. In addition to the log-transformed offset for total nuclei sampled, an offset for target length (gene or domain) was used to normalize for differences in the length of the target gene/domain. A gene-specific dispersion parameter was estimated in the gene-level model via dispformula = ~gene, motivated by diagnostic analyses showing that a single dispersion parameter was insufficient to capture gene-level heterogeneity in mutational burden. Model optimization for the gene-level model was performed using the BFGS algorithm due to convergence failures with the default optimizer, owing to the gene-specific dispersion parameterization. Estimated marginal means per gene and per domain were derived from their respective models using the emmeans package, back-transformed to the response scale at unit offset values to express predictions as mutational burden normalized for total nuclei and target length, and averaged over all other covariates at their observed means.

Separate negative binomial mixed-effects models were fitted for each gene to estimate covariate effects independently, without the constraints of a shared effects structure, incorporating the same fixed- and random-effects covariates and the log-transformed offset as described above, except for the gene identity term. Model optimization was uniformly performed using the BFGS algorithm as the default optimizer encountered convergence issues for *UNC13A*. For each fixed-effect covariate of interest, model coefficients were exponentiated to obtain effect estimates with 95% Wald confidence intervals. *P*-values were Bonferroni-corrected across genes with a significance threshold of $P < 0.05$.

Model fit was assessed using DHARMa scaled residual diagnostics,[12] including Kolmogorov–Smirnov tests for uniformity, dispersion tests and outlier tests. All models converged successfully with no residual overdispersion (all dispersion ratios <1.2, $P$ >0.05) and no evidence of zero-inflation (all DHARMa $P$ >0.05).

### Supplementary figures

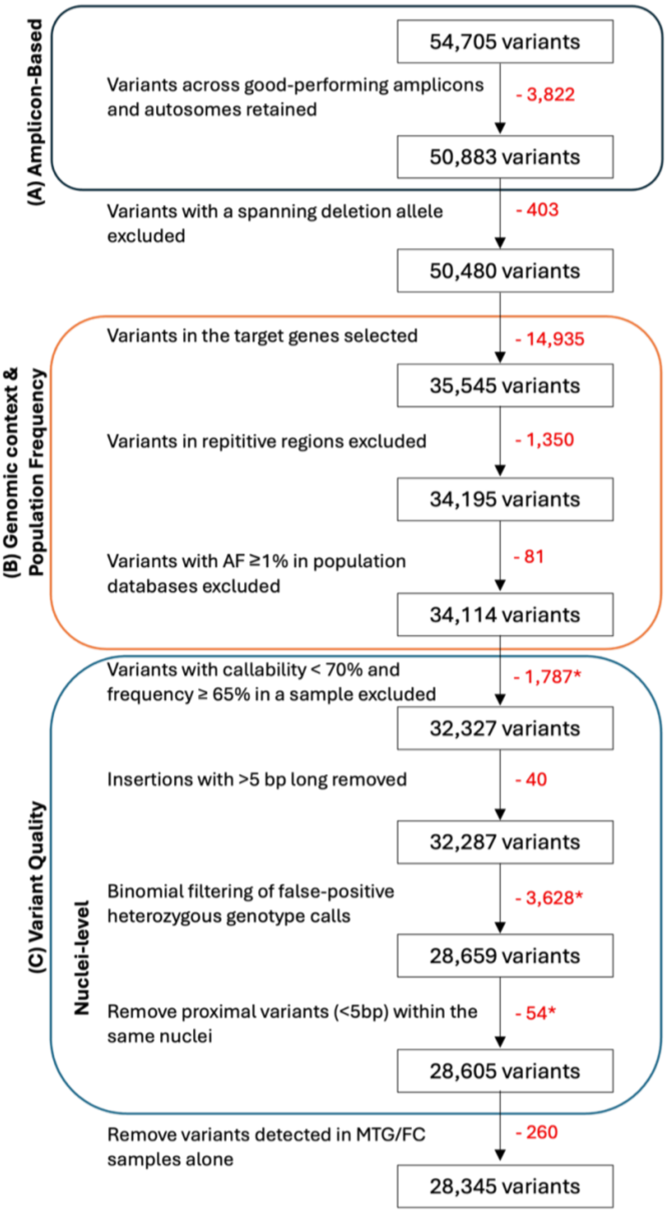


**Supplementary Figure 1. Overview of the variant filtering pipeline.** Summary of the sequential filtering steps applied to remove low-confidence variants. **(A)** Variants mapping to poorly performing amplicons ($n$ = 5) or to sex chromosome-specific amplicons ($n$ = 12) were excluded. Variants with a spanning deletion allele were also removed. **(B)** Variants were further filtered by genomic context and population frequency. Variants mapping to target genes were retained, excluding those associated with multiplexing amplicons. Variants within repetitive regions of the target genes were discarded. In addition, variants with an allele frequency ≥ 1% in gnomAD (exomes or genomes) or in the 1000 Genomes reference panel were excluded. **(C)** To minimize false-positive somatic variant calls, additional filtering criteria were applied. In a sample, variants with callability below 70% (low-confidence variants) and allele frequency ≥ 65% (likely germline variants) were reclassified as missing (genotype = 3) for that sample. Insertions longer than 5 bp were removed. At the single-nucleus level, heterozygous calls with insufficient alternate allele support (one-sided binomial test against $p_0 = 0.5$, Benjamini-Hochberg FDR-corrected $P$-value < 0.05) were reclassified as homozygous reference (genotype = 0). Finally, proximal variants (within 5 bp of another variant in the same nucleus) were set to missing. *Represents somatic variants removed entirely from the dataset.

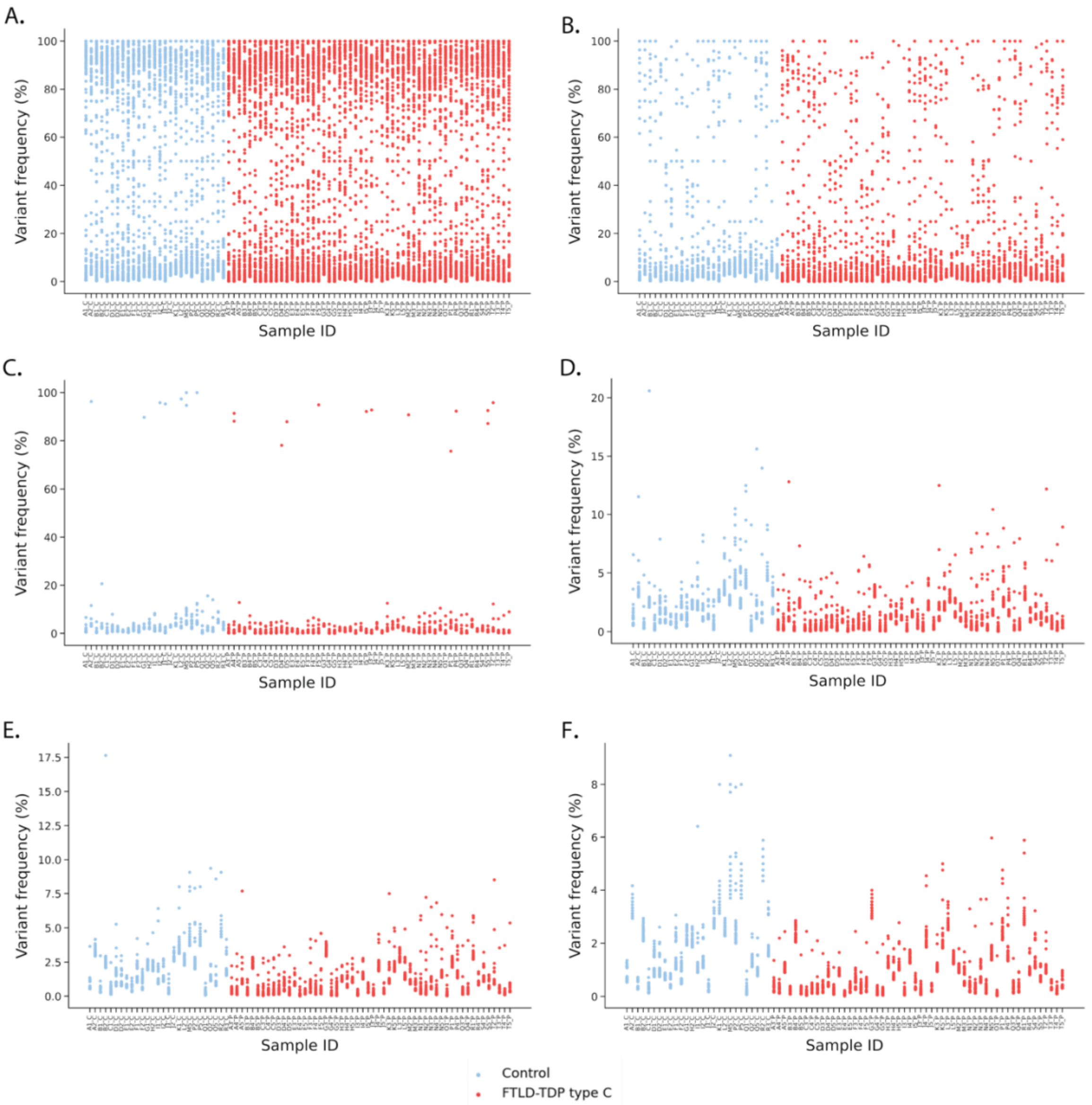


**Supplementary Figure 2. Sequential variant filtering steps and their effects on variant allele frequencies per sample.** Scatter plots showing the allele frequency of individual variants per sample (y-axis) across all samples (x-axis). **(A)** All detected variants prior to filtering. **(B)** Variants remaining after excluding those with an allele frequency ≥ 1% in gnomAD (exomes or genomes) or in the 1000 Genomes reference panel, and those overlapping repetitive genomic elements as annotated by RepeatMasker. **(C)** Variants with callability below 70% at the given genomic position are excluded. **(D)** Variants with allele frequency ≥ 65% are removed as likely germline, based on orthogonal validation against whole-genome sequencing data from 11 shared frontal cortex brain tissue samples. **(E)** Allele frequencies recalculated after binomial threshold filtering of heterozygous calls and exclusion of variants occurring within a 5 bp window in the same nucleus. **(F)** Allele frequencies after retaining coding variants only, corresponding to the variant frequencies shown in **Fig. 1G**.

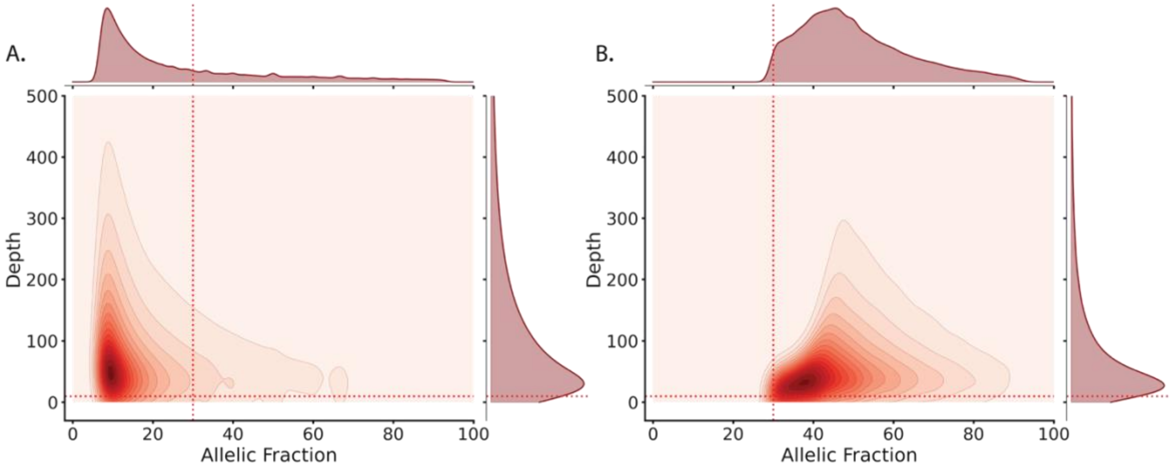


**Supplementary Figure 3. Read depth and variant allele frequency (VAF) for heterozygous variant calls before and after variant filtering.** Read depth versus VAF for heterozygous variant calls depicted (A) prior to filtering (54,705 variants; 5,924,531 heterozygous calls) and (B) following variant quality filtering (28,605 variants; 141,301 heterozygous calls) across 41,128 neuronal nuclei in our cohort. Calls with VAF < 30% or read depth < 10 were converted to missing prior to sequential filtering (**Supplementary Figs. 1** and **2**). Marginal densities are shown for each axis. The y-axis is truncated at 500 for visual clarity.

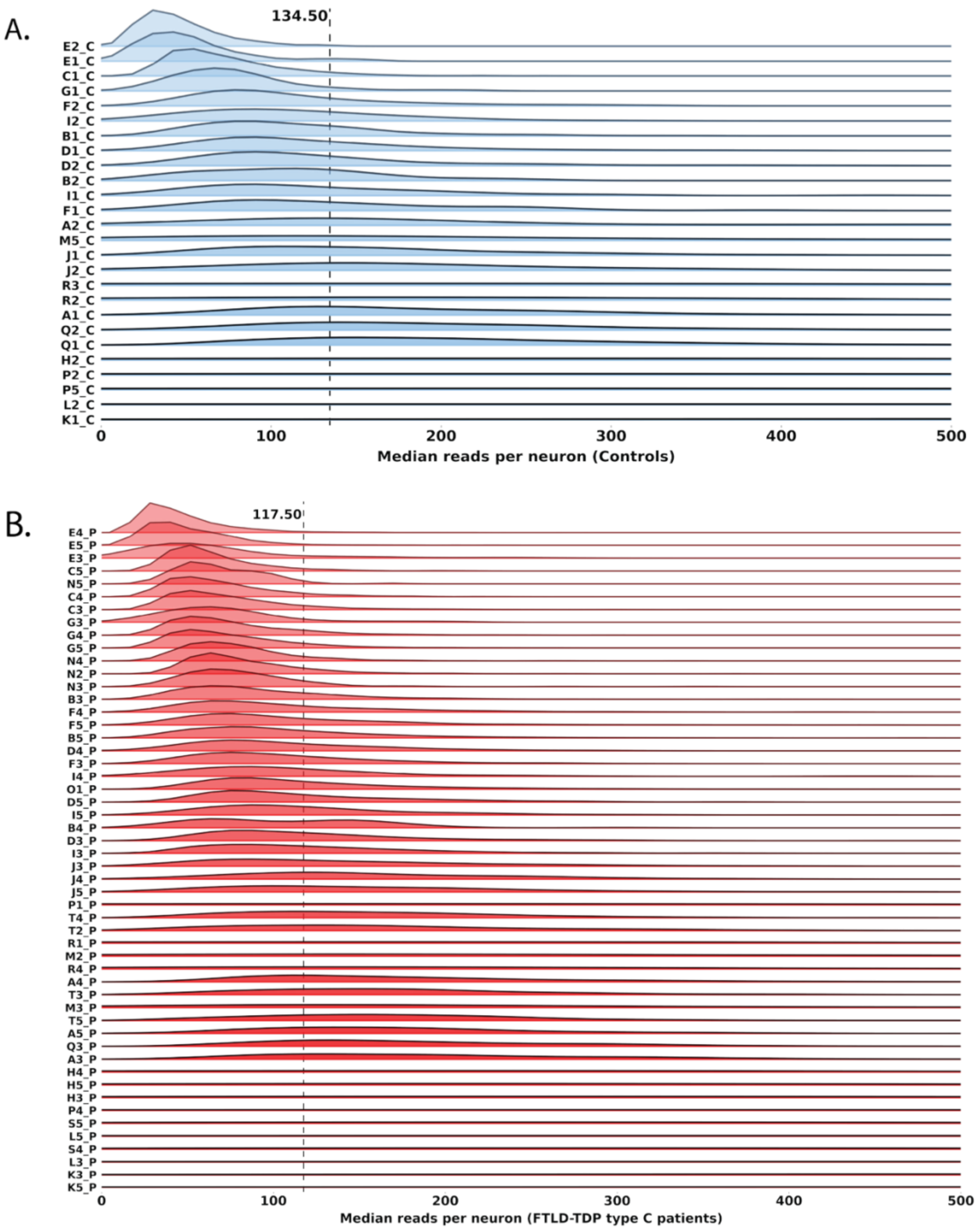


**Supplementary Figure 4. Median read depth across samples.** Distribution of per-base median sequencing depth across individual neuronal nuclei (x-axis) for each superior temporal gyrus sample (y-axis) from **(A)** controls and **(B)** FTLD-TDP type C individuals. The per-nucleus median depth is computed across all coding positions after variant filtering. The vertical dotted line denotes the group-level median depth. Samples are ordered from top to bottom by decreasing total number of captured nuclei; samples with fewer captured nuclei show less dense distributions. The x-axis is truncated at 500 for visual clarity.

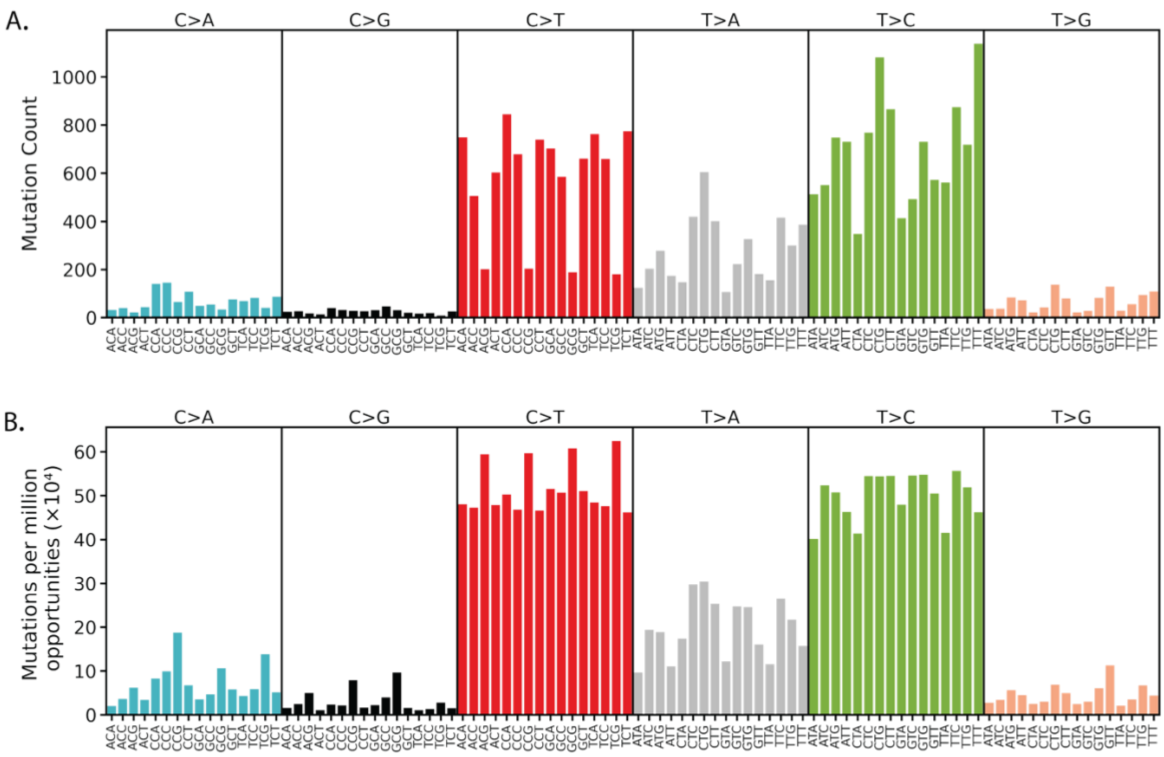


**Supplementary Figure 5. Mutational spectrum of 27,109 somatic single-nucleotide variants (sSNV) detected in our cohort. (A)** Absolute sSNV counts and **(B)** trinucleotide opportunity-normalized sSNV counts across 96 contexts (x-axis; COSMIC conventions). In (B), observed counts were divided by the frequency of each trinucleotide within the sequenced intervals (BED-defined capture regions; GRCh38) and expressed as mutations per million opportunities ($\times 10^4$), correcting for sequence composition bias inherent to amplicon sequencing. Substitution classes are colour-coded, and all variants are shown in pyrimidine-centred notation.

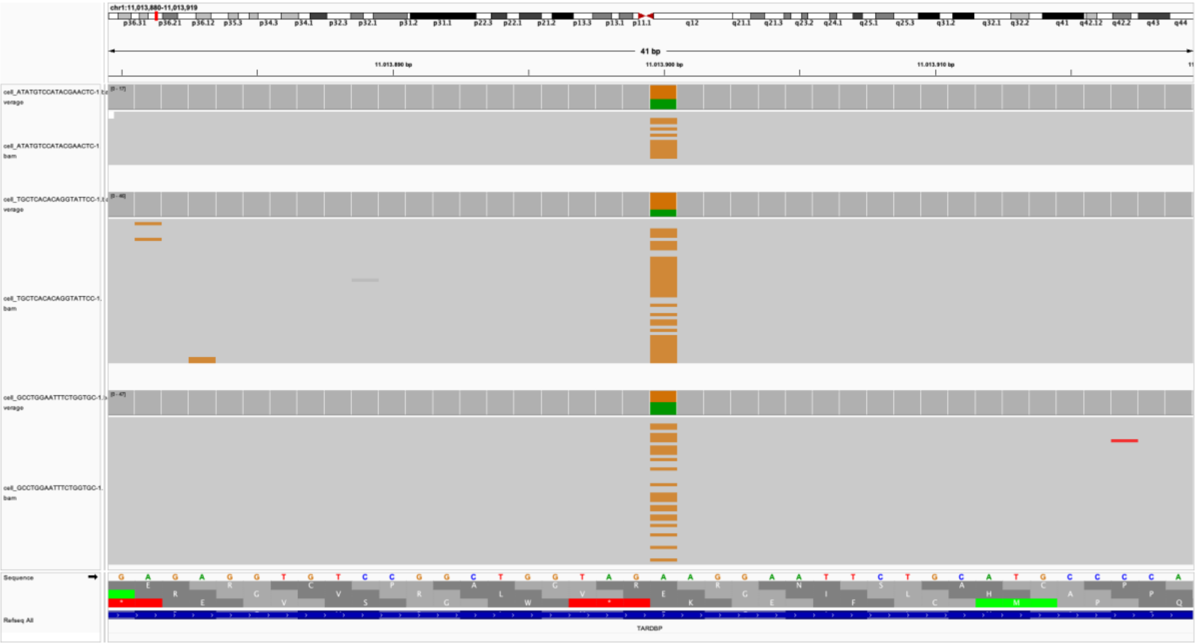

**Supplementary Figure 6. *TARDBP* p.E58G variant in three neuronal nuclei from sample D4_P.** Integrative Genomics Viewer visualization of three neuronal nuclei from patient D4_P, carrying a heterozygous A>G substitution at chr1:11,013,900 (GRCh38), corresponding to the *TARDBP* p.E58G missense variant. Read pileups and coverage tracks are shown for each nucleus across the region chr1:11,013,880–11,013,919, with a maximum observed read depth of 47 reads per nucleus. The variant position is highlighted. The reference genome sequence and RefSeq annotation of *TARDBP* are shown at the bottom.

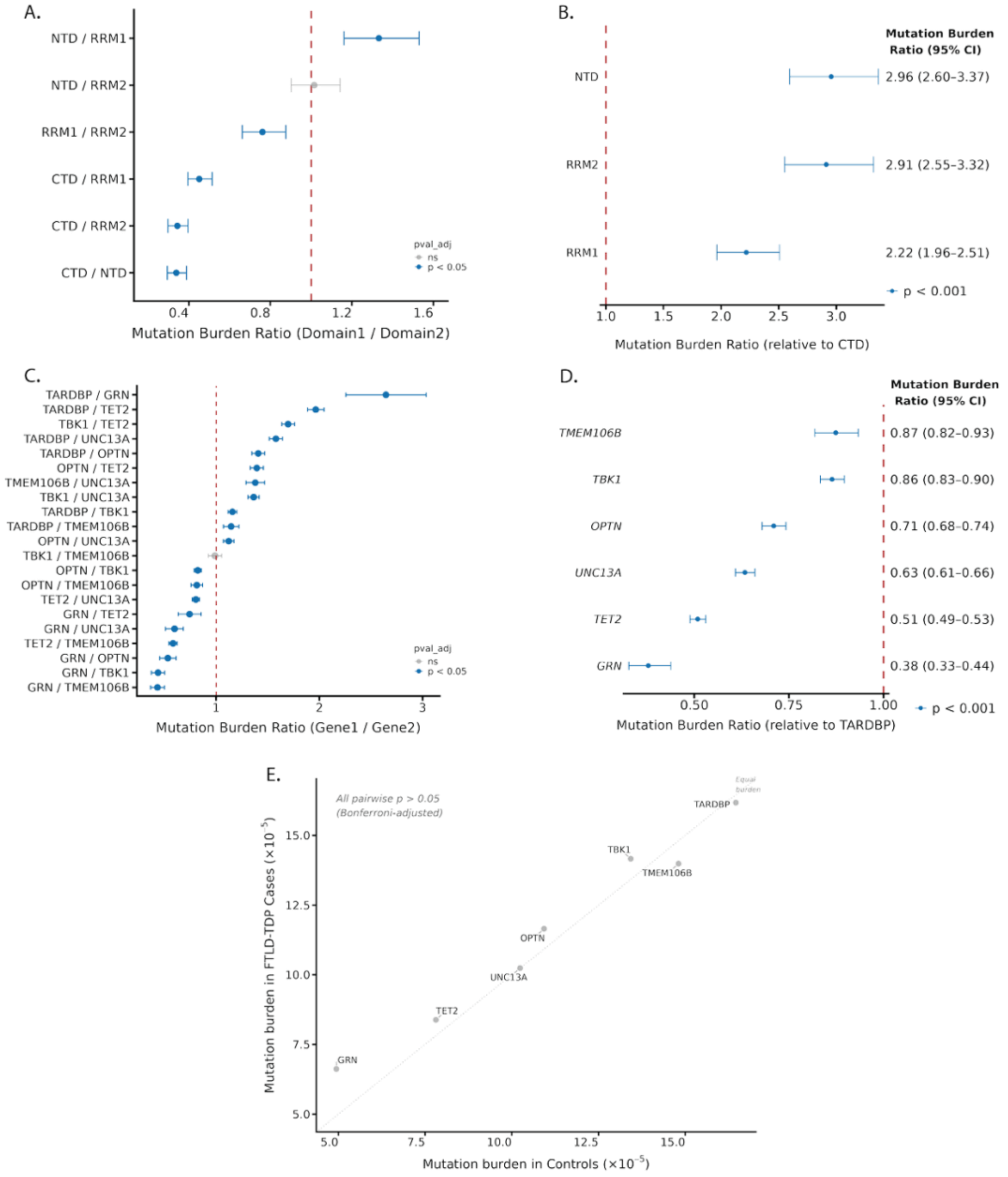


**Supplementary Figure 7. Combined analysis of somatic mutational burden across targeted genes and TDP-43 protein domains. (A–B)** Comparison of mutational burden across TDP-43 protein domains (NTD, RRM1, RRM2, CTD) and **(C-D)** across targeted genes (*TARDBP, TBK1, OPTN, UNC13A, TMEM106B, GRN, TET2*). **(A)** All pairwise mutation burden ratios and **(B)** ratios relative to the CTD, derived from a negative binomial mixed model (see Methods). **(C)** Equivalent pairwise and **(D)** *TARDBP*-referenced analyses across targeted genes. In plots A–D, point estimates and 95% confidence intervals represent mutation burden ratios derived from estimated marginal means at unit cell count and unit sequence length. The vertical dashed line indicates equal mutation burden between the compared domains or genes (ratio = 1). *P*-values were derived from Wald z-tests on the log scale and Bonferroni-adjusted for the number of pairwise tests. Point colour denotes adjusted significance (blue: $p < 0.05$; grey: ns). **(E)** Gene-level, nuclei- and length-normalized mutation burden (estimated marginal means from the gene × Condition model) in FTLD-TDP type C (y-axis) versus controls (x-axis); the dotted line indicates equal burden. Per-gene mutation burden did not differ significantly between conditions after Bonferroni correction across genes ($P > 0.05$).
